# Resistance and resilience of soil microbiomes under climate change

**DOI:** 10.1101/2023.08.04.551981

**Authors:** Julia A. Boyle, Bridget K. Murphy, Ingo Ensminger, John R. Stinchcombe, Megan. E. Frederickson

## Abstract

Soil microbiomes play key roles in plant productivity and nutrient cycling, and we need to understand whether and how they will withstand the effects of global climate change. We exposed *in situ* soil microbial communities to multiple rounds of heat, drought, or both treatments, and profiled microbial communities with 16S rRNA and ITS amplicon sequencing during and after these climatic changes, and then tested how domain and symbiotic lifestyle affected responses. Fungal community composition strongly shifted due to drought and its legacy. In contrast, bacterial community composition resisted change during the experiment, but still was affected by the legacy of drought. We identified fungal and bacterial taxa with differential abundance due to heat and drought and found that taxa affected during climate events are not necessarily the taxa affected in recovery periods, showing the complexity and importance of legacy effects. Additionally, we found evidence that symbiotic groups of microbes important to plant performance respond in diverse ways to climate treatments and their legacy, suggesting plants may be impacted by past climatic events like drought and warming even if they do not experience the event themselves.

## Introduction

Anthropogenic climate change is increasing both global temperature and the frequency and severity of weather events like drought (Thuiller 2007). The microbial communities in soil are among the largest and most diverse on earth (Singh et al. 2009) and their response to climate change will affect elemental cycling, greenhouse gas production, plant-soil interactions, and ecosystem services and agriculture (Dubey et al. 2019; Jansson & Hofmockel 2020; ASM Report 2022). Here, we examine the effect of repeated heat and drought events on a soil microbiome across time, characterizing how fungal and bacterial communities resist, and recover from, heat and drought events.

Resistance refers to how quickly and how much an ecological community responds to shifting environmental conditions, whereas resilience is measured by the speed and degree to which community composition returns to baseline historical conditions (Bissett et al. 2013; Hartmann et al. 2013; Martiny et al. 2017). Low resilience may lead to lasting changes in a community after the disturbance, known as legacy effects. Microbial communities can resist more or less depending on the type and duration of environmental change (Schimel et al. 2007; Zhou et al. 2020), with evidence of microbes being more resistant to short-term than long-term changes (Pec et al. 2021; Seaton et al. 2021). After an environmental shift, microbial communities are often assumed to have high resilience due to short microbial generation times (Allison and Martiny 2008), but empirical evidence is mixed, as described below.

Heat and drought stress can harm the integrity of microbial cells, leading to reduced and differential survival of microbes, and stress can alter microbial activity through modifying gene expression and enzyme production, with knock-on effects on other microbes and the soil environment (Bérard et al. 2015). As a result, the resistance and resilience of microbial communities to environmental change are often characterized by changes in alpha or beta diversity, microbial community composition and microbial biomass (Allison & Treseder 2008; Castro et al. 2010; Sheik et al. 2011; Melillo et al. 2017; Zhou et al. 2020; Pec et al. 2021; Seaton et al. 2021). Empirically, there appears to be little consistent pattern in microbial biomass, richness, and diversity responses to warming or drought. For instance, microbial biomass can increase (Castro et al. 2010) or decrease (Allison & Treseder 2008) in response to heat, with a decrease in microbial biomass accompanying a loss of overall soil carbon (Melillo et al. 2017).

Heat can increase species richness (Allison & Treseder 2008) or decrease it (Sheik et al. 2011), with a meta-analysis of studies finding no overall trend of warming or altered precipitation on microbial alpha diversity (Zhou et al. 2020). The same meta-analysis also found no overall trend of altered precipitation, but a positive effect of warming, on microbial beta diversity (Zhou et al. 2020). Despite the overall lack of consistent patterns, empirical research frequently reports significant changes in microbial relative abundance and identity (community composition) due to heat or drought (Allison & Treseder 2008; Castro et al. 2010; Zhou et al. 2020; Pec et al. 2021). Changes in soil microbial community composition due to climatic events can persist long after the event (Averill et al. 2016; Martiny et al. 2017; de Nijs et al. 2019) or sometimes do not persist (Evans & Wallenstein 2012; Rousk et al. 2013; Dacal et al. 2022), suggesting resistance and resilience may vary amongst soil microbial communities (Waldrop & Firestone 2006). Changes in community composition can be further characterized through time using metrics like turnover (appearances and disappearances) and mean rank shifts (Hallett et al. 2016), where greater turnover and larger mean rank shifts can indicate less resistant and resilient communities.

Resistance and resilience can also be considered from the perspective of continued ecological services (Allison & Martiny 2008), which are less likely to be altered in communities with high stability. Communities are considered stable when groups of taxa are negatively covarying and asynchronous, as decreases in one group can be compensated for by increases in another, potentially recouping total organismal biomass and ecological function (Thibaut & Connolly 2013; Huang et al. 2020; Valencia et al. 2020). Therefore, understanding the effect of environmental stress on microbial community stability is important for predicting changes in ecosystem services (Wagg et al. 2018; Hernandez et al. 2021). In practice, microbial community stability can be estimated with microbial covariances, microbial networks, and by examining microbial synchrony in response to a stressor. Environmental stresses like water availability and lack of nutrients have been shown to increase positive correlations between microbial soil taxa, decreasing community stability and jeopardizing ecological function (Hernandez et al. 2021; Gao et al. 2022); thus we predict a destabilizing effect of heat and drought on the soil microbiome.

Differences in resistance and resilience are likely among component members of the microbiome, due to the vast taxonomic and functional diversity of soil microbes (Singh et al. 2009). A phylogenetic signal in microbial response to climatic events is probable because many microbial traits are phylogenetically conserved, although we lack data on the phylogenetic signal of ecologically relevant phenotypes like heat and drought resistance (Goberna and Verdú 2016). If resistance and resilience are phylogenetically conserved, this lends predictive power to microbial response to climate events. We predict large differences in resistance and resilience between domains, due to their phylogenetic distance and lifestyle differences. For example, bacteria are generally fast-growing, with short generation times and the ability to horizontally transfer potentially adaptive genes (Ochman et al. 2000), while fungi show a myriad of life history strategies, many of which require multiple stages (Andrews 1992) and thus longer generation times. In the soil food web, bacteria are considered a ‘fast’ energy channel that cycles nutrients rapidly, while fungi are considered a ‘slow’ energy channel that cycles nutrients slowly (Rooney et al. 2006; Bardgett & Caruso 2020). Consistent with this classification, turnover times of soil fungi are estimated to be an order of magnitude longer than those of soil bacteria (Rousk & Bååth 2011). Slow fungal turnover times mean that it could take longer to observe a shift in fungal than in bacterial community composition, such that the effect of climate stressors on fungal composition may manifest more slowly, but persist for longer.

Microbial lifestyle (e.g., symbiosis propensity and type) could also have implications for how well a microbe can resist or recover from climatic events. Many soil microbes engage in symbiosis with host organisms, including plants, animals, and macroscopic fungi (Berendsen et al. 2012), with the outcome of symbiosis ranging from negative to positive for the host. One major group of plant symbionts is rhizobia, a polyphyletic group of soil bacteria that facultatively nodulate legume roots, where they exchange fixed nitrogen for fixed carbon (Beringer et al. 1979). We highlight rhizobia because their status as mutualistic symbionts of legumes is well-characterized; furthermore, they are critical to the global nitrogen cycle and modern agricultural practices (Hirsch & Mauchline 2015). Rhizobia genomes are large for bacteria and contain plasmids that encode genes required for nodule formation and nitrogen fixation (MacLean et al. 2007), and this extra genetic material could hamper growth rates and select for genomic streamlining under warming (Hessen et al. 2010; Sabath et al. 2013; Simonsen 2021). Some parasites may be encumbered by similar problems, for example, the bacterial plant pathogen *Agrobacterium tumefaciens* also carries large plasmids that contain the tumor-inducing genes required for its lifestyle (White & Winans 2007), which could slow replication times. In addition, plants have a ubiquitous, often mutualistic, symbiosis with mycorrhizal fungi where mycorrhizae provide mineral nutrients to plants in return for carbon and improve plant tolerance of water stress by extending the root system (Kakouridis et al. 2022). However, mycorrhizal fungi are often obligately symbiotic (Smith & Read 1997), and thus the fate of a mycorrhizal fungus under climate stress may depend on the performance of their plant partner and vice versa, to positive or negative effect (Stachowicz 2001; Kiers et al. 2010). In sum, there are clear pathways through which the life history and symbiotic nature of a microbe could underlie facets of the microbe’s resistance or resilience to climate events.

More empirical evidence on the response of microbial communities to climate events is needed to understand the existing mixed results and to determine what factors underpin microbial resistance and resilience (Jansson & Hofmockel 2020). We conducted a factorial experiment to examine the response of soil microbial communities to the interacting climate factors of heat and drought, with an emphasis on measuring resiliency and legacy effects. We identified microbial groups and lifestyles particularly altered under climate change and discuss why they may be differentially affected. Based on the findings from the literature described above, we hypothesized that fungal communities would be more resistant but less resilient than bacterial communities to climatic perturbations due to their slower growth and turnover times. For similar reasons, we hypothesized that symbiotic microbes would be more resistant but less resilient than the microbial community at large, with the strongest effects on microbes that lack a partner in the experimental plots, because they are in an ecological context without the benefits of symbiosis, but where they bear many of the costs. Given that microbes are a major contributor to global nutrient cycling (ASM Report 2022), interact strongly with species in higher trophic levels (Berendsen et al. 2012), and contribute to ecosystem functioning (Ochoa-Hueso et al. 2020), microbial resistance and resilience to climate change will have far-reaching consequences.

## Methods

### Site description and sampling

We conducted this study at the Koffler Scientific Reserve (KSR, www.ksr.utoronto.ca) in Ontario, Canada (latitude 44°01’48”N, longitude 79°32’01”W) in an old field environment (Figure 1). The average annual precipitation is 767 mm, with the average high temperature reaching 27°C in July and the average low temperature reaching −11 in January (Koffler Scientific Reserve 2023).

**Figure 1.**
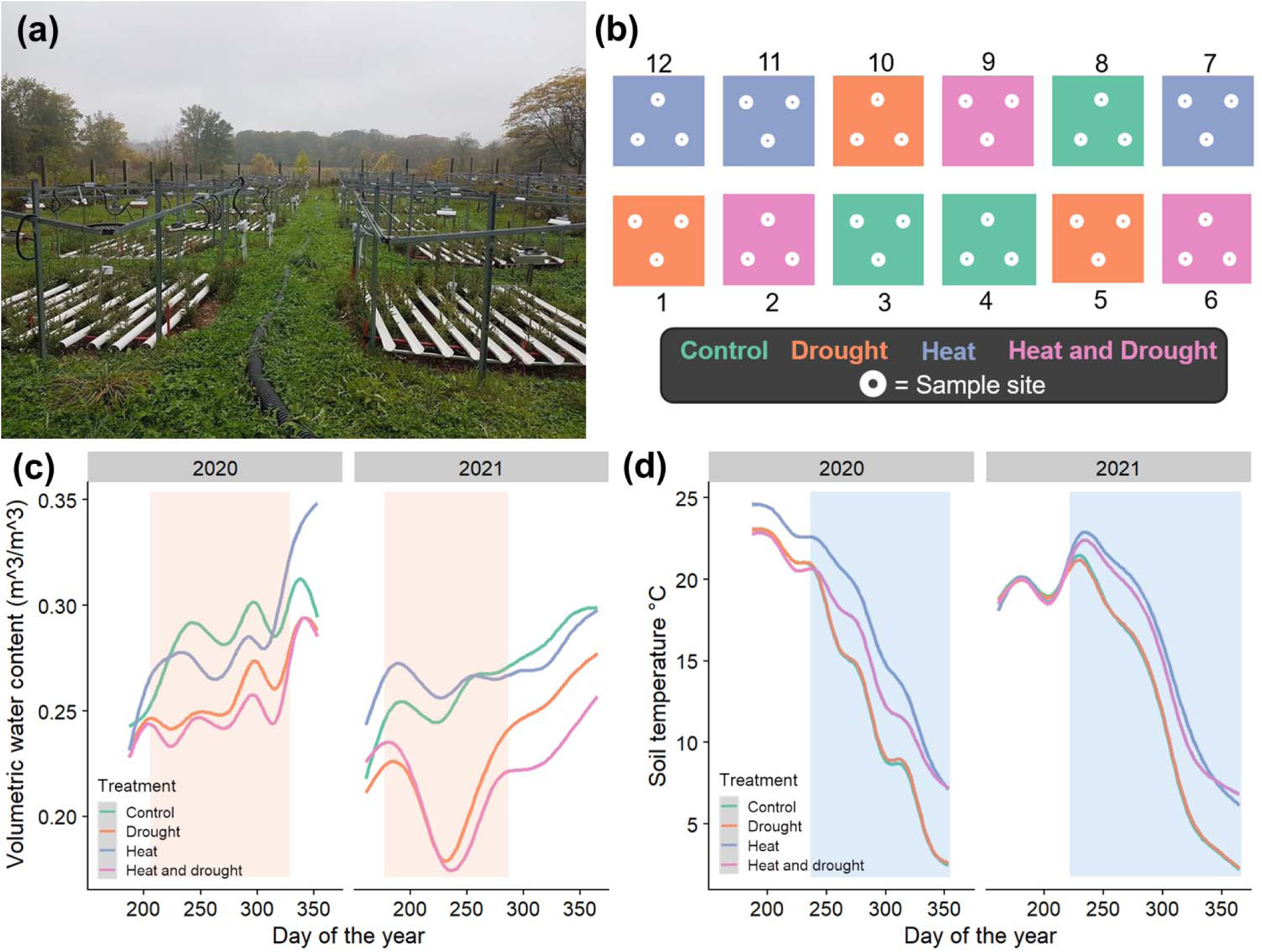
Experimental design at Koffler Scientific Reserve. (a) The temperature-free-air-controlled enhancement plots showing the infrared heaters and rainout structures, as well as the water drainage system. Photo by Julia Boyle. (b) Schematic diagram of the experimental plots (numbered) and their climate treatments. Dots represent sampled sites within plots. (c) Drought treatment was effective at reducing soil volumetric water content. (d) Heat treatment was effective at increasing soil temperature. Lines in (c) and (d) show GAM smoothed means of treatments and overlays indicate when drought (orange) or heat (blue) was actively applied.

The temperature-free-air-controlled enhancement experiment at KSR was established following Kimball et al. (2008). Plots are 3m in diameter, with six plots having si× 100W infrared heaters each (Mor Electric Heating Association) and six plots having equivalent dummy infrared heaters. In 2017, we excavated twelve plots 30 cm deep and filled them with a 1:1:1 mixture of local soil, peat moss, and sand; then, we planted white spruce seedlings (*Picea glauca,* provided by Natural Resources Canada, Laurentian Forestry Centre, Quebec QC) in the plots. A top layer of commercially supplied cedar bark mulch was added in 2020 and 2021, and weeding occurred biweekly to keep plant cover constant. Starting in 2020, we applied climate treatments to plots: control, heat, drought, or heat and drought (Figure 1). Heat and drought treatments were established to study the physiological and molecular responses of different white spruce genotypes to climatic stressors through the growing season and into early winter; those results will be presented elsewhere. The heaters ran day and night in the heated plots, while the drought plots had concave-up (U) slotted rainout structures that covered ∼50% of the plot surface area and drained rain out and away from all plots. To control for the effects of the structures themselves, non-drought plots had the same rainout structures except in the concave-down position (⋂), allowing all water to pass off the structure and into the plot. There were two periods of active treatment (Appendix 1: Table S1): rainout structures were present July-November 2020, and June-October 2021, for a total of 8 months; heaters were active in heated plots August-December 2020, and August 2021-January 2022, for a total of 9 months. To determine how treatments affected soil conditions, soil volumetric water content (VWC) and temperature were monitored using soil sensors and dataloggers (detailed methods in Appendix 2). During active treatment, the mean soil temperature of heated plots was 3.7 or 3.6 hotter than un-heated plots in 2020 and 2021 respectively (Figure 1). The mean soil VWC during droughts was 0.28 (m^3^/m^3^) in non-drought plots and 0.25 (m^3^/m^3^) in drought plots in 2020, and in 2021, mean VWC was 0.26 (m^3^/m^3^) in non-drought plots and 0.21 (m^3^/m^3^) in drought plots (Figure 1). So, soil VWC in drought-treated plots was relatively decreased by 10% in 2020 and 20% in 2021.

On June 11th 2021, September 14th 2021, and June 5th 2022, we sampled from the first 10 cm of bulk topsoil from three locations ∼50 cm apart in each plot. We collected the June 2021 and 2022 samples when there were no active climate treatments being applied (plots in a recovery period) to test microbial resilience, and the September 2021 samples when climate treatments were being actively imposed to test resistance. After each collection, samples were stored at −20 until DNA extraction.

### Sequencing and QIIME2

We extracted soil microbial DNA using the QIAGEN DNeasy PowerSoil Pro Kit following the kit protocol and sequenced samples at Genome Quebec (Montréal, Canada) using Illumina MiSeq PE 250bp 16S rRNA amplicon sequencing on the conserved hypervariable V4 region (primer pair 515F-806R) and ITS region (primer pair ITS1FP2-58A2RP3). Samples from 2021 and 2022 were sequenced separately. We used Quantitative Insights Into Microbial Ecology 2 (QIIME2) v.2022.2 (Bolyen et al. 2019) to trim sequences and then denoised sequences with DADA2 (Callahan et al. 2016) to obtain amplicon sequence variants (ASVs). Using QIIME2, we removed ASVs that had fewer than 10 reads across all samples, and assigned taxonomy using the ‘sklearn’ feature classifier (Pedregosa et al. 2011); we used Greengenes 16S V4 region reference for bacteria (McDonald et al. 2012), and UNITE (Nilsson et al. 2019) version 9.0 with dynamic clustering of global and 97% singletons for fungi. We then filtered out reads assigned as cyanobacteria and mitochondria to remove plant and animal DNA. Finally, for each of the bacterial datasets we constructed a phylogeny using QIIME2’s MAFFT (Katoh & Standley 2013) and FastTree (Price et al. 2010) functions to obtain a rooted tree.

### Statistical Analysis

We performed analyses in R v4.2.0 (R Core Team 2022), with the ‘tidyverse’ (Wickham et al. 2019), ‘phyloseq’ (McMurdie & Holmes 2013), and ‘microbiome’ (Lahti & Shetty 2019) packages. For statistical tests, we merged the subsamples of plots such that each plot had one set of reads for each time point and domain, then rarefied samples to the minimum read number for that sampling time point (Appendix 1: Table S2). We analyzed the data from each time point separately unless otherwise stated. Models followed the same general structure, with heat treatment, drought treatment, and the heat × drought interaction as predictors, unless otherwise stated. To visualize communities, we used principal coordinates analysis (PCoA) on the rarefied relative abundance data, using weighted UniFrac distance for bacteria and Bray-Curtis distance for fungi. Next, we tested for differences in rarefied relative community composition using adonis2 permutational analysis of variance with the Bray-Curtis method and 9999 permutations (‘vegan’ package, Oksanen et al. 2022); results did not change based on the order of factors. We used betadisper (‘vegan’ package, Oksanen et al. 2022) and the base R dist function to measure the distance between centroids when heat or drought was applied, which provides a quantitative effect size of treatment on community composition.

We leveraged the repeated sampling of our climate-treated plots to understand how the soil microbiome changed over time. We tested for differences in observed ASV richness, evenness, Shannon diversity, and Simpson’s diversity across timepoints, using the rarefied datasets and linear models, then used type 3 ANOVAs from the ‘car’ package (Fox & Weisburg 2019) on the linear models to calculate F values. We measured temporal diversity indices using the ‘codyn’ package (Hallett et al. 2020), including the mean rank order shifts of genera and turnover (total, appearances, and disappearances) within communities of the same climate treatments. We also used ‘codyn’ to measure two community stability metrics, synchrony and variance ratio, with ASVs aggregated to genus. Synchrony compares the variance of aggregated genera abundances with the summed variance of individual genera (Loreau & de Mazancourt 2008), with 0 indicating complete asynchrony, and 1 indicating complete synchrony. The variance ratio measures the pairwise covariance of genera, where a value of 1 indicates no covariance, a value below 1 indicates negative covariance, and a value above 1 indicate positive covariance (Schluter 1984); we permuted the data 999 times to generate a null estimate with confidence intervals, and an observed variance ratio that was higher or lower than the confidence interval indicated a significantly positive or negative covariance among genera, respectively. We tested for differences in synchrony, rank shifts, and turnover using linear models in the same way as for differences in the alpha diversity metrics, except the rank shifts and turnover models included time comparison as an added interacting predictor. To further compare pairwise correlations of genera across treatment and domain during active treatment, we combined fungal and bacterial datasets aggregated at the genus level and used SparCC implemented through the ‘Spiec-Easi’ package (Friedman & Alm 2012; Kurtz et al. 2015) to create co-occurrence networks. We permuted the correlations 100 times in each network to create a null expectation for significance testing, and adjusted *p* values using the false discovery rate. Then, to determine how correlated genera are with each other and if groups of genera correlate together more in each network, we determined the Kleinberg’s hub centrality scores of genera, the number of clusters, and modularity.

To detect differentially abundant taxa, we used the analysis of compositions of microbiomes with bias correction (‘ANCOMBC’) package (Lin & Peddada 2020), which identifies significantly differentially abundant taxa while accounting for bias from library size. We performed this analysis at the genus level with Holm’s method p-value adjustment. We visualized the magnitude and overlap of significant log fold responses using heat maps and ‘UpSetR’ (Conway et al. 2017). Then, we tested for consistent differences in the response of resistant or resilient microbes by using student’s *t*-tests to compare the log fold response of genera in active treatment to genera in recovery periods. We tested domains separately and their responses to heat and drought separately, and used natural log transformed responses as needed to improve normality. Next, we determined whether genera correlated to many other genera were less resistant to climate events by using a MANOVA with log fold response in active heat and drought as the responses, and hub score in active control conditions as the predictor.

To test whether bacterial genera’s response to heat and drought is phylogenetically conserved, we used the log fold change in abundance due to heat or drought calculated from ANCOMBC as continuous traits. Using phylosig from the ‘phytools’ package (Revell 2024) we calculated Pagel’s λ (Pagel 1999) for both traits at each timepoint separately, where a value of 0 indicates phylogenetic independence (no significant signal) and a value of 1 indicates traits follow the expected distribution under Brownian motion (Münkemüller et al. 2012). We tested for phylogenetic signal only in our bacterial data, as fungal phylogenetic reconstructions using the ITS region are not reliable. Then, we visualized log fold change due to heat and drought along the bacterial phylogeny using the ‘phylosignal’ package (Keck et al. 2016).

To assess treatment effects on groups of known symbiotic bacteria, we leveraged catalogues by Li et al. (2023) and Lajudie et al. (2019) to create separate subset datasets of genera containing plant beneficial bacteria (PBB), phytopathogens, and rhizobia. We tested for differences in diversity and community composition as described above using the rarefied datasets aggregated at genus-level. In our experimental plots, rhizobia responded to climate events without any legume partners, due to weeding. Next, we assessed treatment effects in symbiotic fungi. As white spruce was the focal plant host in our plots, we subset the data to families containing ectomycorrhizal fungi that commonly associate with white spruce (Lazaruk et al. 2008; Smith et al. 2015; Nadeau & Khasa 2016) and tested for differences in their relative abundance using linear models. We identified more symbiotic fungi by classifying fungi into guilds using FUNGuild annotation (Nguyen et al. 2016) and kept classifications only when ‘Probable’ or ‘Highly Probable.’ We concatenated classifications across years and relativized each guild’s raw read number to sample read number, then used repeated measures ANOVAs with time included as a factor and plot as the error term. The response variables were the relative abundance of symbiotic guilds of interest: parasites, pathogens, endophytes, ectomycorrhiza, non-ectomycorrhizal mycorrhizae (e.g. ericoid, arbuscular), and all mycorrhizae. We also tested the relative abundance of the saprotroph guild for comparison. Response variables were natural log transformed as needed to improve the normality of error distributions.

## Results

### Fungal communities

#### Fungal community composition and diversity were not resistant or resilient to drought

During active treatment, fungal community composition was not resistant; it was strongly shaped by drought (*F_1,8_*=1.24, *p*<0.05; Appendix S1: Table S4), but not by heat or heat × drought. Significant drought, but not heat or heat × drought, effects on fungal community composition also persisted to the second recovery period (drought *F_1,8_*=2.35, *p*<0.05; Figure 2B, Appendix S1: Table S4), meaning fungal communities were not resilient and slow to recover after experiencing climate treatments.

**Figure 2.**
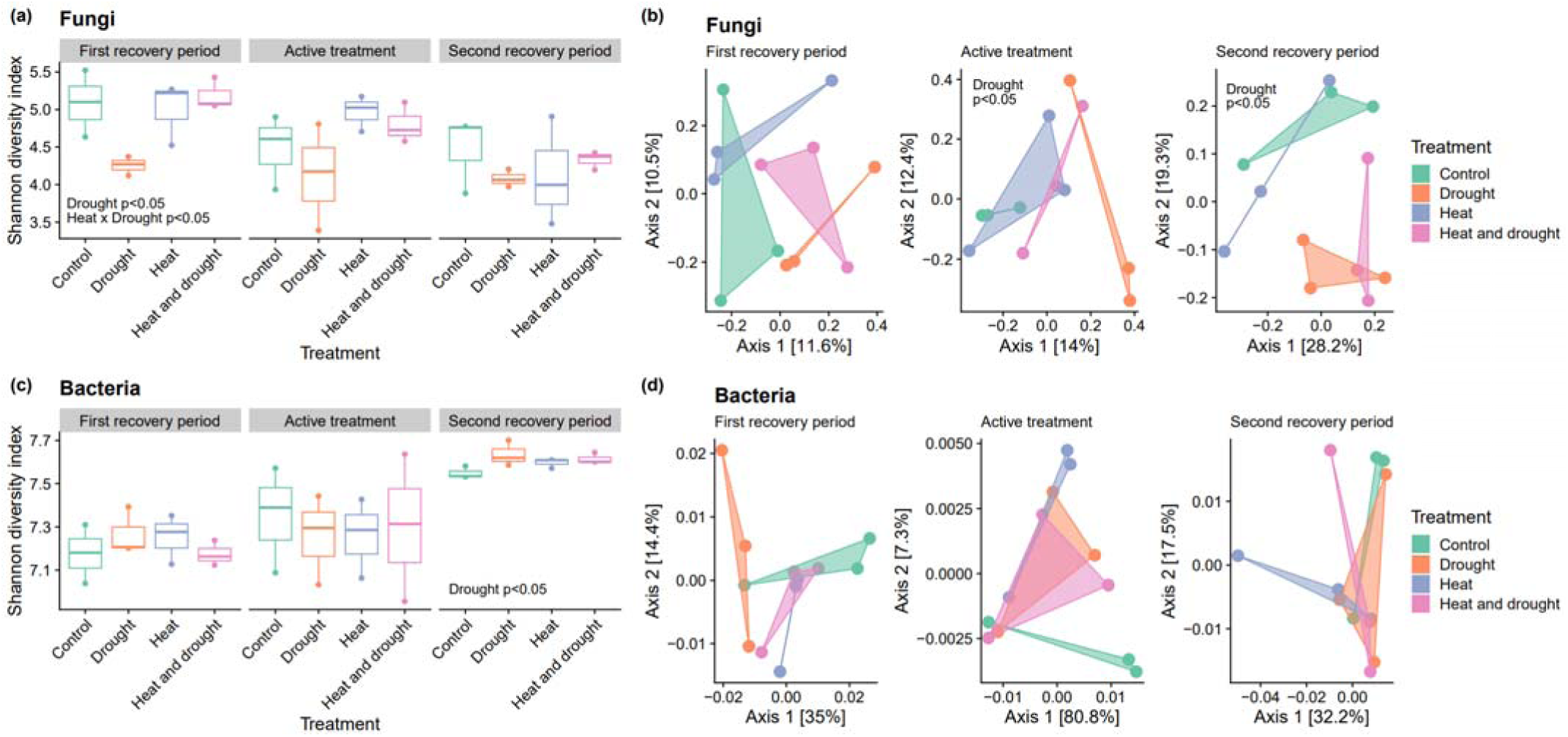
Community structure of fungal and bacterial soil microbes across climate conditions and time. Boxplots of Shannon diversity across time in fungal (a) and bacterial (c) microbial communities. PCoA plots show fungal Bray-Curtis dissimilarity (b) and bacterial weighted Unifrac distances (d) of the microbial communities. Only significant effects of heat, drought, and their interaction are displayed on the panels.

Drought-induced shifts in community composition during active treatment did not change the alpha diversity of fungal communities; Shannon diversity, Simpson’s diversity, evenness, and observed richness were not significantly affected by any active experimental treatment (Appendix S1: Table S3). The only statistically significant effects of climate treatments on alpha diversity occurred in the first recovery period, when legacy of drought caused a 16% reduction in Shannon diversity (*F_1,8_*=9.67, *p*<0.05; Appendix S1: Table S3) and a 4% reduction in Simpson’s diversity compared to control plots (*F_1,8_*=8.74, *p*<0.05; Appendix S1: Table S3). Heat mitigated the effect of drought on fungal alpha diversity in the first recovery period, significantly so for Shannon diversity (*F_1,8_*=7.19, *p*<0.05; Figure 2A; Appendix S1: Table S3) and marginally significantly for Simpson’s diversity (*F_1,8_*=5.04, *p*=0.055; Appendix S1: Table S3). Drought and heat × drought effects on Shannon diversity were driven by significant changes in evenness (*F_1,8_>*7.01, *p*<0.05 for all effects; Appendix S1: Table S3), not by changes in observed species richness (Appendix S1: Table S3).

Underlying community-wide shifts, we identified 161 genera (full list in Figure 4 and Figure 5) that shifted in estimated absolute abundance in response to heat, drought, or both. There were only 30 genera that significantly responded to both active treatment and treatment legacies (Figure 4, Figure5, Appendix S3: Figure S4). Drought and its legacy increased and decreased the estimated absolute abundance of fungal genera to similar extents during treatment and recovery periods (*t*=1.21, *df* =87.2, *p*=0.231; Figure 4, Appendix S3: Figure S5). In contrast, active heat decreased the estimated absolute abundance of fungal genera while heat legacy overall increased estimated abundance; the distribution of fungal responses to heat was significantly different between active and recovery periods (*t*=-3.01, *df* =15.5, *p*<0.01; Figure 5, Appendix S3: Figure S5). *Saccharomycetales* (yeasts) showed strikingly higher relative abundance in drought plots (Figure 3), driven by the *Babjevia* genus (∼3x log fold increase in drought, Holm’s adjusted *p*<0.05; Figure 4). We also saw interactions of fungal phenology and climate treatments. For example, the average relative abundance of *Pezizales* in the control plots increased by 92% from June to September 2021, then decreased 83% between September 2021 and June 2022 (Figure 3), suggesting *Pezizales* are more naturally abundant in September. In contrast, the relative abundance of *Pezizales* in drought-treated plots increased by an average of only 77% between June and September 2021, and decreased 86% between September 2021 and June 2022. Drought dampened the phenological increase in *Pezizales* abundance, and as a result *Pezizales* had one of the largest negative log fold changes in response to drought among fungi (Holm’s adjusted *p*<0.05; Figure 4).

**Figure 3.**
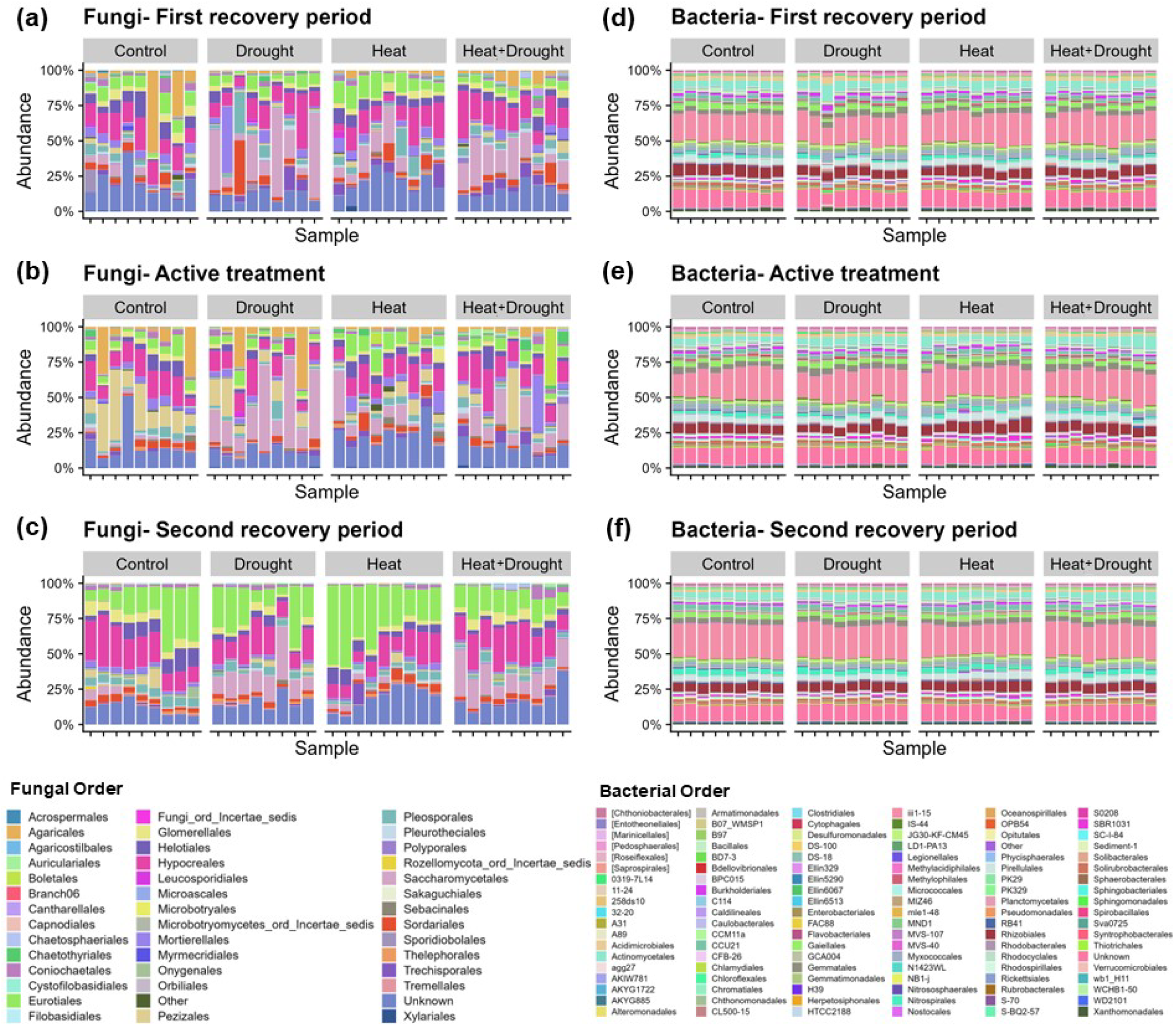
Community composition over time for each subsample in each treatment. (a)-(c) Fungi relative abundances at the order level. (d)-(f) Bacteria relative abundance at the order level. Orders were aggregated and included when they were at least 1% of the compositional abundance, and found in at least 30% of samples. Plot order within treatment groups remains consistent, with 2-3 samples per plot.

**Figure 4.**
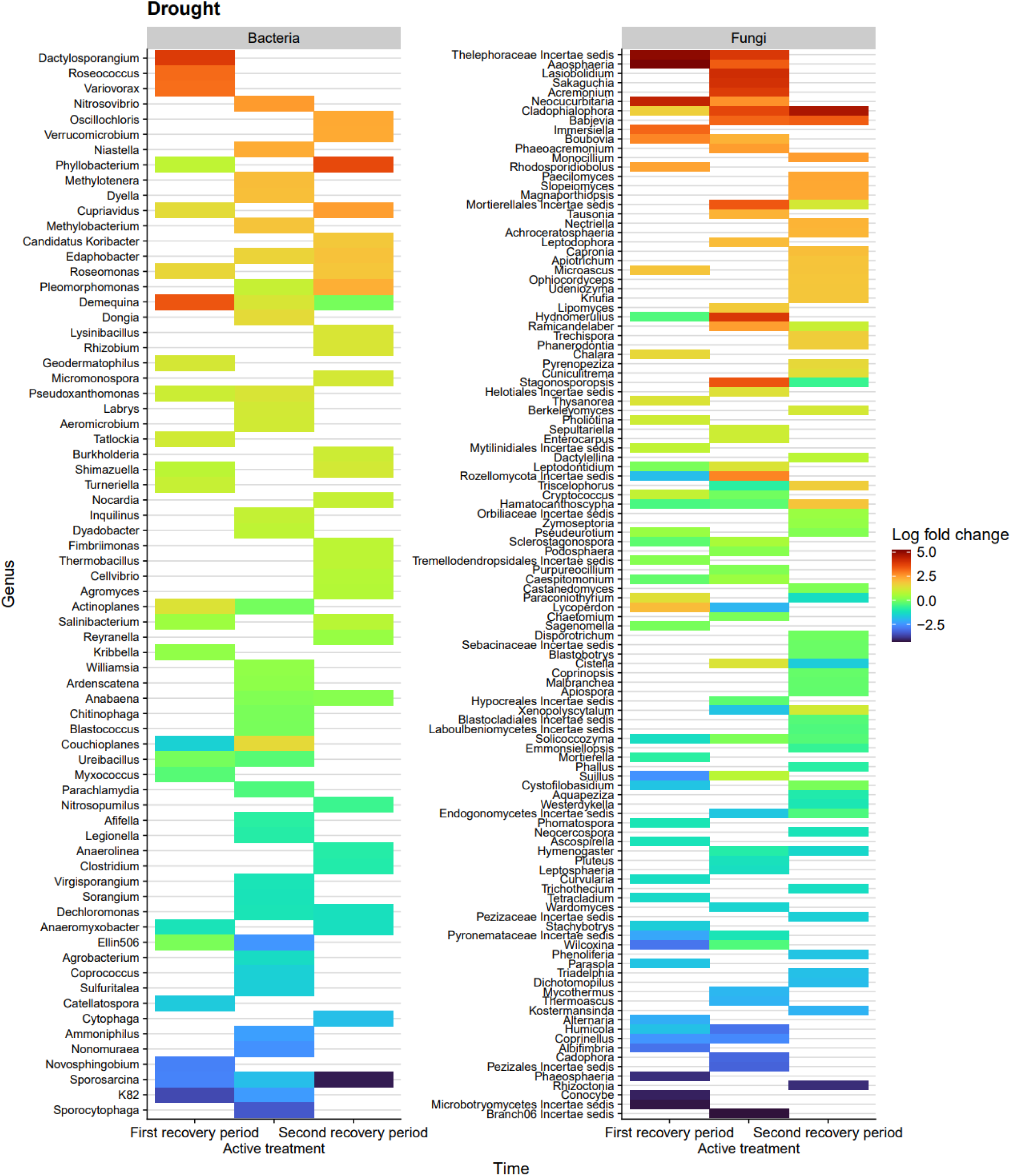
Differentially abundant microbe genera due to drought across all sampling times, compared to control plots. Taxa shown are significant (p<0.05) with Holm’s method *p*-value adjustment, grouped by domain, and ordered by descending average log fold change.

**Figure 5.**
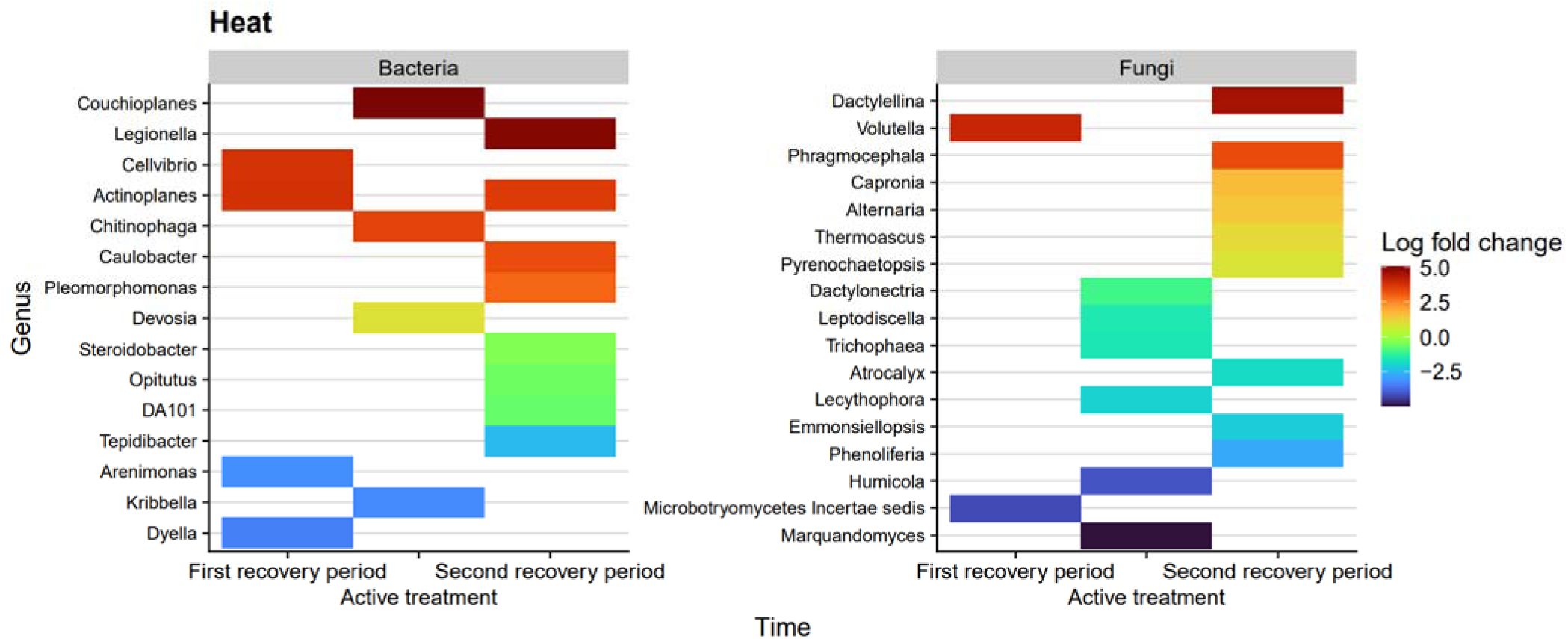
Differentially abundant microbe genera due to heat across all sampling times, compared to control plots. Taxa shown are significant (p<0.05) with Holm’s method *p*-value adjustment, grouped by domain, and ordered by descending average log fold change.

#### Heat altered fungal community dynamics across time

We measured community change through time within treatments and found evidence of low fungal resistance and resilience. First, we tested the effect of heat and drought on turnover in genera across timepoints. Many genera were present across all time points (n=193), with the most unique genera occurring in the second recovery period (n=75) (Appendix S3: Figure S1). All aspects of turnover significantly depended on which time was being compared to the first recovery period (*F_1,8_*>10.1, *p*<0.01 for all effects, Appendix S1: Table S5; Appendix S3: Figure S2), for example, there were 42% more disappearances that occurred between the first recovery period and active treatment compared to the disappearances between recovery periods. Heat significantly altered genera appearances and disappearances (*F_1,8_*>6.70, *p*<0.05 for all effects, Appendix S1:Table S5), but not drought. The effect of heat differed depending on which periods were compared (*F_1,8_>*7.22, *p*<0.05 for heat × time, Appendix S1:Table S5). Between first recovery period and active treatment, active heat alone increased the appearances of new genera by 25% and decreased disappearances by 23%. In contrast, between recovery periods, heat decreased new appearances by 17% while disappearances increased 40% (*F_1,8_*>6.70, *p*<0.05 for all effects, Appendix S1:Table S5). Total turnover between recovery periods was not significant (Appendix S1:Table S5). Climate treatment and time comparison did not significantly alter the average relative change in species rank abundance (Appendix S1: Table S5; Appendix S3: Figure S2).

We next considered how climate treatments affected community stability, e.g. whether the abundance of taxa changed at the same time and in the same direction in response to climate treatments. The fungal genera in control plots positively covaried, but not significantly so, with a variance ratio of 1.55; heat alone and drought alone increased the variance ratio by 0.5 each, and their effects were synergistic in the heat and drought treatment (variance ratio of 2.69), indicating climate treatments increased the strength of positive covariance among genera. Variance ratios were only significantly higher than permuted estimates when heat was applied. Overall synchrony of fungi was low, ranging between 0.1-0.3, and the difference among treatments was not statistically significant (Appendix S1: Table S5, Appendix S3: Figure S2).

#### Mutualistic and parasitic fungal guilds differed in their resistance and resilience

Our analysis of symbiotic fungal guilds showed the relative abundance of parasitic fungal guilds (parasitic on other fungi, lichen, plants, or insects) increased 36% in response to drought (*F_1,8_*=13.0, *p*<0.01; Figure 6, Appendix S1: Table S6). The guilds of pathogens, endophytes, and saprotrophs were not significantly affected by the climate treatments (Appendix S1: Table S6). The mycorrhizal guild as a whole significantly decreased by an average of 59% in relative abundance due to heat (*F_1,8_*=6.15, *p*<0.05; Appendix S1: Table S6) and had a non-significant trend of 51% lower abundance under drought. Mycorrhizal relative abundance was much greater during the September samples compared to June samples (*F_2,22_*=13.2, *p*<0.001; Appendix S1: Table S6), and 93.4% of mycorrhizal reads in September were ectomycorrhizal, which is not surprising given that the spruce saplings in the plots are ectomycorrhizal. Ectomycorrhizal fungi on their own decreased an average of 61% in heat-treated plots and 50% in drought-treated plots, however the reduction in abundance in response to heat was only marginally significant (*F_1,8_*=5.26, *p*=0.051; Appendix S1: Table S6), likely due to low statistical power. The remaining mycorrhizal reads consisted of ericoid and arbuscular mycorrhiza; on their own, these fungi had 43% and 60% reduced relative abundance due to heat and drought, respectively, throughout all timepoints, although there were no significant differences from control plots (*F_1,8_*<1.70, *p*>0.22 for all effects; Appendix 1: Table S6, Appendix S3: Figure S3). Most mycorrhizal reads were in the families *Pyronemataceae, Herpotrichiellaceae, Ceratobasidiaceae, Entolomataceae, Helotiaceae, Pezizaceae, Thelephoraceae, Myxotrichaceae, Hymenogastraceae,* and *Suillaceae*.

**Figure 6.**
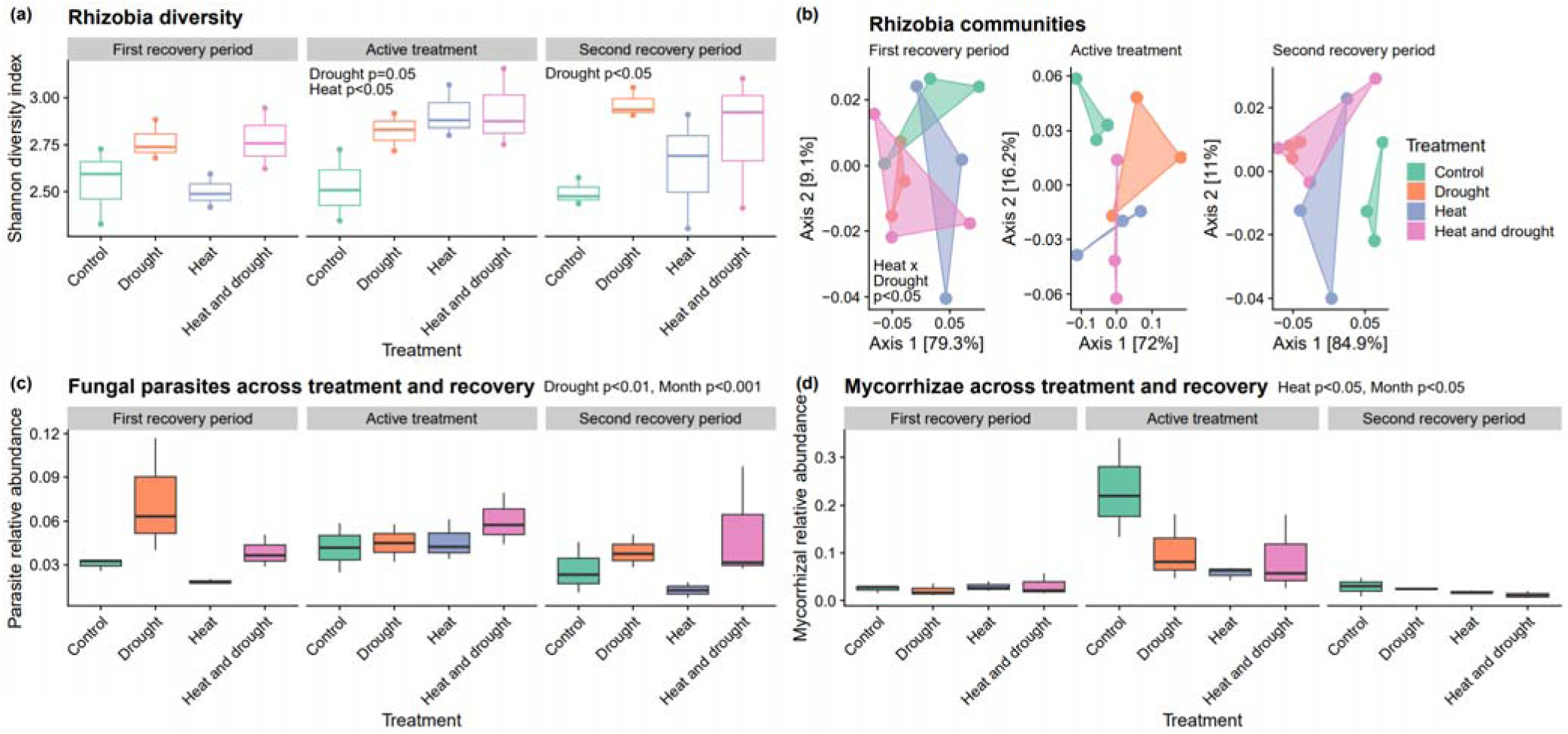
Symbiotic microbes under climate treatments. (a) Rhizobia Shannon diversity. (b) Rhizobia community weighted Unifrac PCoAs. (c) and (d) Fungal parasite and mycorrhizae relative abundance across treatment and time. Only significant main effects of heat, drought, and their interaction are displayed on the panels.

We also examined only ectomycorrhizal fungi known to associate with white spruce. We found that their relative abundance decreased by an average of 58% due to heat and 49% due to drought across all timepoints, similar to the trends for all mycorrhizae. During active treatment, heat significantly reduced white spruce ectomycorrhizal relative abundance in the bulk soil (*F_1,8_*=6.36, *p*<0.05; Appendix 1: Table S7), but the reduction was not significant in recovery periods (*F_1,8_*<3.52, *p*>0.05 for all effects; Appendix 1: Table S7). We identified two differentially abundant ectomycorrhizal genera that associate with *Pinaceae* tree saplings: *Wilcoxina,* and *Suillus* (Dahlberg & Finlay 1999; Yu et al. 2001). *Suillus* and *Wilcoxina* had ∼2.5x and ∼3x log fold decreased estimated absolute abundance due to legacy of drought, but differing responses during active drought, with decreased *Wilcoxina* and increased *Suillus* (Holm’s adjusted *p*<0.05; Figure 4). The *Thelephoraceae* family has an ectomycorrhizal genus that associates with white spruce, *Thelephora* (Lazaruk et al. 2008); interestingly, an unidentified genus from this family was extremely differentially abundant, with a significant ∼5x log fold increase due to drought in the first recovery period, and significant increase during active treatment (Holm’s adjusted *p*<0.05; Figure 4).

### Bacterial communities

#### Bacterial community composition and diversity were resistant and resilient to heat and drought

Bacteria communities were more resistant to the climate events than fungi, with no significant shifts due to active treatments (Appendix S1: Table S8 and 10-13% less distance between centroids during active treatment compared to fungi. Bacteria community composition was also very resilient to climate treatments, showing no lasting compositional changes in recovery periods.

Bacteria diversity was resistant and resilient to climate treatments, especially heat. Shannon diversity, Simpson’s diversity, evenness, and observed richness were not affected during active treatment or the first recovery period (Appendix S1: Table S9). In the second recovery period, the legacy of drought significantly increased bacterial Shannon diversity, but by only 1% (*F_1,8_=*9.30, *p*<0.05; Figure 2C, Appendix S1: Table S9), potentially because of a marginally significant increase in observed richness amounting to an average of 293 ASVs (*F_1,8_=*5.11, *p*=0.054; Appendix S1: Table S9). No other alpha diversity metrics were affected in the second recovery period (Appendix S1: Table S8-S9).

Despite whole bacterial communities having strong resistance to climate treatments, we identified 102 differentially abundant bacterial genera due to heat and drought during active treatment and recovery periods (Holm’s adjusted *p*<0.05; Figure 4, Figure 5). Only 16 taxa were significantly affected during both active treatment and recovery periods (Appendix 3: Figure S4). Bacteria significantly affected by heat and heat legacy tended to increase in estimated absolute abundance (Figure 5), and there was no significant difference in the log_10_ transformed distribution of responses between active and recovery periods (*t*=0.295, *df*=4.53, *p*=0.781; Appendix S3: Figure S5). Drought and drought legacy increased the estimated absolute abundance of some bacterial taxa, and decreased the abundance of others, but the distribution of effects was not significantly different in active and recovery periods (*t*=-1.32, *df*=25.4, *p*=0.198; Appendix S3: Figure S5). We detected significant phylogenetic signal in bacterial response in only the second recovery period to both heat (Pagel’s λ=0.31, *p*<0.001; Appendix S1: Table S11, Appendix S3: Figure S6) and drought (Pagel’s λ=0.19, *p*<0.01; Appendix S1: Table S11, Appendix S3: Figure S7). This phylogenetic signal may in part be driven by genera belonging to the orders *Bacillales* and *Clostridiales* in the phylum *Firmicutes*; the *Firmicutes* clade showed consistent decreases in log-fold estimated absolute abundance to both heat and drought throughout the experiment, but especially in the second recovery period (Appendix S3: Figure S4-S5).

#### Bacterial community dynamics were not affected by heat or drought

Bacterial community dynamics did not differ significantly in response to treatment, again suggesting high resistance and resilience. Most bacterial genera were present across all sampling times (n=167), and active treatment had the most unique bacterial genera (n=35) (Appendix S3: Figure S1). There were no significant differences in the mean rank shift or turnover of bacterial genera due to either climate treatments, nor were there significant differences between treatment and recovery periods (Appendix S1: Table S10, Appendix S3: Figure S2). Though there was no effect of climate, bacterial genera had very high synchrony ranging between 0.84-0.96, and genera in all climate-treated communities were significantly positively covarying with a variance ratio of ∼1.40 (Appendix S3: Figure S2).

#### Heat and drought more strongly affected symbiotic bacterial guilds compared to all bacteria

Phytopathogenic bacteria had high resistance and resilience to climate treatments, though they were more affected by treatment compared to the whole bacterial community. There was no significant effect of climate treatment on phytopathogenic alpha diversity metrics or community composition in the first recovery period or active treatment (Appendix S1: Table S12-S13). In the second recovery period, heat significantly decreased bacterial phytopathogen evenness by an average of 2% (*F_1,8_*=11.8, *p*≤0.01, Appendix S1: Table S12), and drought significantly shifted community composition (*F_1,8_*=1.75, *p*≤0.05, Appendix S1: Table S13). The ANCOM-BC differential abundance analysis found the estimated absolute abundance of a potentially phytopathogenic genus *Agrobacterium* decreased 1.3x log fold due to drought, and a human pathogen, *Legionella*, significantly increased 4.9x log fold due to the legacy of heat (Figure 4).

The plant beneficial bacteria (PBB) showed high resistance but low resilience to drought. In the first recovery period and active treatment, there were no significant changes to PBB’s alpha diversity metrics or community composition (Appendix S1: Table S14-S15). However, in the second recovery period drought significantly increased PBB’s Shannon and Simpson diversity by an average of 2% each (*F_1,8_*>11.6, p≤0.01 for all, Appendix S1: Table S14). This increase in diversity was driven by significantly higher observed ASV richness in drought; drought-only plots increased by a mean 47 ASVs compared to control plots, but heat significantly dampened this increase as indicated by the significant heat × drought interaction (*F_1,8_*>8.94, p≤0.05 for all, Appendix S1: Table S14). Community composition of PBB was not significantly changed in the second recovery period (Appendix S1: Table S15). The ANCOM-BC differential abundance analysis found that active drought and legacy of drought increased plant growth-promoting rhizobacteria in the soil (Figure 4), for example *Variovorax* spp. increased by 3x log fold, and *Pseudoxanthomonas* spp. by 1-1.3x log fold, which can help plants tolerate abiotic stresses such as drought (Kang et al. 2014; Kaushal 2019; Singh et al. 2021).

Rhizobia showed strong community-wide responses both during and after the heat and drought treatments. During active treatment, heat and drought significantly increased the rhizobia guild’s Shannon diversity by an average of 13% and significantly increased Simpson’s diversity by an average of 4% (*F_1,8_*>5.29, p≤0.05 for all, Appendix S1: Table S16). Heat significantly increased observed richness by an average of 32% (an additional 8 ASVs) during active treatment (*F_1,8_*=6.58, p<0.05, Appendix S1: Table S16). Evenness and community composition were not significantly changed during active treatment (Appendix S1: Table S16-S17). In the first recovery period, the rhizobia community composition was significantly predicted by the heat × drought interaction (*F_1,8_*=1.78, p<0.05, Appendix S1: Table S17, Figure 6), but no alpha diversity metrics were affected (Appendix S1: Table S16). In the second recovery period, Shannon diversity and Simpson’s diversity remained significantly increased due to legacy of drought by 14% and 5% respectively (*F_1,8_*>5.46, p<0.05 for all, Appendix S1: Table S16, Figure 6), and there was a marginally significant increase in evenness (*F_1,8_*=5.22, p=0.052; Appendix S1: Table S16). Community composition was not significantly affected by heat and drought in the second recovery period (Appendix S1: Table S17). The ANCOM-BC differential abundance analysis showed that drought and legacy of drought increased the log fold abundances of many rhizobia genera (*Burkholderia* 1x, *Cupriavidus* 1.4-2.4x*, Methylobacterium* 1.9x*, Rhizobium* 1.2x*, Phyllobacterium* 0.87-3.6x).

### Heat and drought altered associations in microbial community networks

We analyzed how co-occurrence networks that included both fungi and bacteria were altered by heat and drought during active treatment. The networks indicate groups of taxa with correlated abundances, potentially because of species interactions or niche similarity. Each network, regardless of climate treatment, formed three clusters, and the global clustering coefficients ranged between 0.75-0.78 (Appendix S1: Table S18). Modular networks indicate distinct groupings of taxa that are highly correlated among themselves and less correlated with taxa in other groupings. Modularity increased with both heat and drought, starting at 0.35 in control conditions and peaking at 0.41 when both heat and drought were applied (Appendix S1: Table S18). We identified 37 significant correlations between pairs of genera in control conditions, of which 22% were positive correlations; 60% of the significant correlations in control conditions were between fungi and bacteria (Appendix S1: Table S18). The heat treatment had 43 significant correlations total, but the number of fungi-fungi interactions doubled compared to control conditions (Appendix S1: Table S18). Supporting the variance ratio analyses, heat tended to shift correlations to be more positive: 49% of correlations were positive in the heat treatment. Drought dramatically decreased the number of significant correlations. In the drought treatment, there was only one significant positive correlation between fungi and bacteria, while in the heat and drought treatment there were only 9 significant correlations total (Appendix S1: Table S18). Significantly correlated pairs of microbes are provided as a supplemental file.

Microbes correlated to many other microbes, e.g. hub taxa, were different between each network (Appendix S1: Table S18). The average hub score decreased when drought or heat was applied, compared to control plots, however microbes with a low hub score in control conditions tended to remain the same or increase their score with heat and drought, while microbes with a high hub score in control conditions tended to remain the same or decrease their hub scores in heat and drought (Appendix S3: Figure S8). The hub score of a genus in control conditions significantly predicted how that genus responded to drought (*F_1,59_*=9.19, p<0.01, Appendix S1: Table S19), where genera correlated to many other genera decreased more in active drought. Response to heat was not significantly predicted by hub score (*F_1,59_*=3.31, p=0.07, Appendix S1: Table S19).

## Discussion

Our results suggest that climate change, and especially drought, will have more persistent effects on soil fungi than on soil bacteria. In fungal communities, drought shaped community composition, legacy of drought decreased diversity, and heat and drought altered the relative abundance of symbiotic fungi important to plant communities. Heat changed the strength and sign of covariance of fungal genera and increased temporal dissimilarity within heated communities. In contrast, bacterial communities were largely resistant to climate treatments, although legacy of drought tended to increase diversity and drought and drought legacy increased the relative abundance of some plant beneficial bacteria. We discuss the implications of the differences between these members of the microbial community, and the effects of climatic events on symbiotic microbes below.

### Fungi are less resistant and resilient than bacteria

Microbial communities experienced legacy effects from the climate treatments, with the effect of drought being stronger than the effect of heat. Our study site regularly experiences extreme annual swings in temperature, potentially conditioning microbes to resist and recover faster from heat (Hawkes & Keitt 2015). In contrast to some studies (de Vries et al. 2018; Zhou et al. 2020; Pec et al. 2021), even relatively short-term climate treatments had strong immediate and lasting impacts on fungal communities; droughted fungal communities did not resemble control communities during or after treatment. Moreover, fungal community resilience was lower than bacterial resilience to drought, as indicated by the significant compositional changes in the second recovery period. The other climate factor, heat, decreased the stability of fungal communities by increasing the positive covariance between genera, potentially changing future resistance and resilience to climate events. The differences in response between fungal and bacterial communities suggest there is phylogenetic conservation of ecologically relevant traits at the domain level and given that we detected phylogenetic signal in response to heat and drought at the genus level in bacteria, ecologically relevant traits appear to be conserved at smaller phylogenetic scales as well. Our prediction that fungi would be more resistant but less resilient to climate change than bacteria was only partially supported; their more complex life history strategies and longer generation times may contribute to fungi’s lack of resilience under climate stress, but do not prevent immediate shifts in relative abundance or presence.

In contrast to our prediction, the bacterial community composition and dynamics in our plots were very resistant to climate treatments, despite literature predicting high turnover rates (Rousk & Bååth 2011) that could allow quick responses. These results are consistent with climate conditions affecting slow-growing microbes more, and fungal communities having a higher proportion of very slow-growing microbes compared to bacteria, thus showing greater change. While some literature suggests slow-growing microbes have lower relative fitness under more extreme climate conditions (Sabath et al. 2013), slow-growing microbes may also benefit from climate change relative to faster-growing microbes, especially if heat and drought make resources scarce or reduce the cost of slower growth (Konstantinidis & Tiedje 2004; Abreu et al. 2023).

The life histories of bacteria and fungi likely affected their resistance and resilience to climatic shifts, although differences in dispersal could have also played a role. It is possible that climate treatments may have influenced local bacterial populations, but dispersal was high enough among bacterial metacommunities to mask these effects. If bacteria could disperse easily between our plots, their community compositions would converge more readily and the strength of signal would be smaller (Gianuca et al. 2017), while if fungi cannot disperse as easily, shifts in composition would be more evident. Actual rates and distances of microbial dispersal are not well described, but there is evidence that fungi are more dispersal-limited than bacteria (Zhang et al. 2021) meaning fungi may be less able to escape a poor environment. While we cannot rule out the role of dispersal, dispersal did not mask the significant effects of treatment on rhizobia and other specific bacterial genera. While surprising, highly resistant bacterial communities are not uncommon; for example, Waring and Hawkes (2018) also observed no direct compositional changes to altered precipitation. Despite no significant changes in community composition, the bacteria in our plots likely experienced physiological changes and costs (Schimel et al. 2007; Evans & Wallenstein 2013; Allison & Goulden 2017) as well as turnover in allele frequencies and gene functions (Chase et al. 2021) that we could not detect with 16S rRNA sequencing, which may have then contributed to drought’s legacy effects on bacterial diversity.

The community-wide shifts in composition and diversity we detected are likely a product of both the direct effects of heat and drought on microbes and the indirect effects of climate treatment on microbes mediated through species interactions, such that microbes not directly affected by climate may still have increased or decreased in relative abundance (Bardgett & Caruso 2020). The increase in positive correlations between genera due to heat suggest that climate stress can reveal axes of niche similarity among microbes that are hidden under less stressful conditions. Our results align with new research suggesting that abiotic stress destabilizes microbial networks through increased positive correlations (Hernandez et al. 2021; Gao et al. 2022). Furthermore, our results suggest that these correlations between genera can affect microbial resistance and resilience, given that highly connected hub taxa were more negatively affected by active drought. While network analyses do not directly measure microbe-microbe interactions, they do reflect the net pattern of how microbes interact with each other, respond to environmental variation, and indirect effects. Indirect effects of climate can also be mediated through plant-microbe or animal-microbe interactions, where stressed organisms may alter the soil environment in some way, for example plants may secrete different root exudates (Karlowsky et al. 2018; Bakhshandeh et al. 2019), or stressed organisms may themselves be an altered environment for symbionts due to factors like growth or immune responses (Nejat & Mantri 2017). While examining differential abundance trends over time, it became apparent that microbes affected during active treatment were not necessarily the ones affected in recovery periods (Figure 4, Figure 5 Appendix S3: Figure S4). Sensitive fungi showed especially different responses to heat in active and recovery periods, with the sign of mean response changing (Appendix S3: Figure S5). These observations demonstrate the complexity of legacy effects and resiliency and suggests a strong role of persisting direct and indirect effects of climate on microbial communities.

The fungal community, composed of predominantly saprotrophs and mycorrhiza, showed low resistance and resilience indicating that the ecosystem services they provide may change under future climate conditions. Fungi are major decomposers of organic material, and therefore unlock a large supply of nutrients for higher soil trophic levels (Boddy & Watkinson 1995). Mycorrhizal fungi contribute to nutrient cycling and mineral weathering (Courty et al. 2010), as well as carbon sequestration (Clemmenson et al. 2013). Despite their importance to global elemental cycling and other ecosystem processes, most research on fungal stress responses focuses solely on yeasts (Branco et al 2022). In this experiment, yeasts were common in drought plots, but they did not represent the average fungal response to treatments. Although yeasts thrived under drought, overall fungal diversity was lower after drought and mycorrhizal abundance was reduced after heat, suggesting the breadth of ecosystem services provided by fungal communities will be decreased under future climatic conditions (Mori et al. 2016). Additionally, low resilience in soil communities means that plants that recruit into a population following a climatic shift may interact with an altered pool of microbes even if they do not experience the climate event themselves. Soil microbiomes affect plant performance and fitness (Berendsen et al. 2012; Rubin et al. 2018; Lu et al. 2018; Fitzpatrick et al. 2019) through mechanisms such as nutrient provisioning (Beringer et al. 1979) and altering plant gene expression profiles (Lu et al. 2018). For plants, associations with microbes can result in improved stress tolerance (Gehring et al. 2017; Allsup et al. 2023) or even broadened geographic range size (Afkhami et al. 2014), which can be adaptive under climate change. Therefore, soil microbes modulate how plants respond to climate events (Classen et al. 2015; Rudgers et al. 2020) and soil legacy effects may have far-reaching ecological and environmental consequences.

### Symbiotic microbes respond differently to heat and drought compared to all microbes

Free-living and symbiotic microorganisms may respond differently to climate change, for example if living inside host cells or tissue buffers microorganisms against changes to the abiotic environment (Gulzar et al. 2020). Multiple groups of plant symbionts sometimes responded differently to climate treatments than the average response of bacterial and fungal communities, and ectomycorrhizal fungi that associate with white spruce did not have a universally larger or smaller response to heat and drought compared to other symbiotic microbes. We do not attribute these results to a shifting plant community, because it was held constant throughout the experiment, however results may be influenced by shifts in plant traits under different climate treatments.

Symbiotic bacterial communities were more altered by heat and drought and their legacies compared to the whole bacteria community. Phytopathogens responded to both heat and drought legacies and had a whole community shift, while plant beneficial bacteria diversity increased by multiple metrics due to drought legacy. Li et al. (2023) found that environmental filtering is a large factor structuring the distribution of PBBs across a landscape, and here we provide experimental evidence supporting PBB’s sensitivity to climate and future climate change. The implications of PBB’s higher diversity and a shifted phytopathogen community after drought on plant performance could be an interesting area of future research. The rhizobia guild showed a similar post-drought response compared to the overall bacteria community with an increase in diversity; however, because we also saw significantly higher diversity when heat and drought were being actively applied, this indicates rhizobia positively responded more to active treatment than the overall community and even other symbiotic microbes. Since rhizobia fix biological nitrogen through legume symbiosis (Beringer et al. 1979), their high abundance bodes well for global nitrogen cycles under climate change. This was unexpected, as we predicted rhizobia in particular would be a worse soil competitor under climate treatments due to their relatively large genomes and thus longer replication times. However, if slower-growing microbes are more sensitive to changes in climate, as we saw with our domain comparison, then this rhizobia result is consistent. At higher temperatures, slow-growing marine bacteria cells retain more normal membrane functions and have a lower mortality burden than fast growers (Schaum & Collins 2014; Abreu et al. 2023), which may apply to slow-growing soil bacteria too. Given our results, we might predict the likelihood of symbiosis between specific legume-rhizobia partners and plant-bacteria partners more broadly to differ in future climate conditions. A change in the availability of microbial partners matters because plant-microbe interactions vary in specificity, especially in the legume-rhizobia system (Chen et al. 2021), which could lead to shifts in plant communities and plant ranges (Harrison et al. 2018). The increased abundance and diversity of rhizobia could also shape evolution in the rhizobia-legume system through increased bacterial competition (Burghardt & diCenzo 2023).

While beneficial bacteria increased with our climate treatments, mutualistic fungi were negatively affected. Mycorrhizae, mainly composed of ectomycorrhiza, had significantly lower relative abundance in heated plots, and ectomycorrhiza that associate with the host plant found in the plots did not have a different magnitude or direction of change from mycorrhizae as a whole. Our results suggest mycorrhizae were less resistant to heat than the fungal community overall, which showed no major compositional changes to heat. We also observed significant decreases in the abundance of specific ectomycorrhizal genera in the bulk soil due to active drought and its legacy, and possibly as a result of this decreased abundance, drought has been shown to reduce ectomycorrhizal root colonization (Erlandson et al. 2022). A recent study by Fu et al. (2022) found that drought lowered the diversity of field arbuscular mycorrhizae, and they suggested this shift was mediated by plant communities. In our experiment, because the ectomycorrhiza could associate with the spruce saplings in our plots, it is likely there were negative indirect climate effects on the mycorrhizae mediated through changes in the physiology of the spruce, indirect microbe-microbe interactions, as well as the direct effects of climate treatment. Ectomycorrhizal fungi have a narrow climatic tolerance and thus are predicted to be disproportionately affected by climate change (Baldrian et al. 2023), which our results support.

Mutualistic and parasitic fungi both derive nutrients from hosts, however while heat decreased mycorrhizae, drought increased parasites in recovery periods. If parasitic fungal species become more prevalent in post-drought conditions this could increase the likelihood of soil dwelling organisms hosting a parasite. Drought and heat may be especially problematic in temperate forests, where most trees are ectomycorrhizal (Baldrian et al. 2023) since trees may simultaneously lack their main mutualist partners and experience increased parasite pressure. Organisms hosting parasites may have reduced growth and fitness, leading to reduced ecosystem productivity. Additionally, parasitized hosts may experience more negative effects from climate events than non-parasitized hosts (Baldrian et al. 2023). In positive feedback, pathogens and parasitic microbes are predicted to increase in heated environments in part due to increased susceptibility of stressed hosts (Pritchard 2011; Altizer et al. 2013; Baker et al. 2018). While we saw limited evidence of heat differentially affecting parasites, our results did suggest drought may also play a role in moderating parasite abundances under future climate conditions.

## Conclusions

Soil microbiomes are tightly linked to global nutrient and chemical cycling, and there is a rich literature describing how microbes buffer plants against the abiotic stresses of climate change (e.g. Yang et al. 2009; Lau & Lennon 2012; Rudgers et al. 2020; Marro et al. 2022; Allsup et al. 2023). Given the soil microbiome’s important role in ecosystems, it is important to understand how microbial populations respond to climate events and the factors underlying variation in resistance and resilience. We found that fungi were less resistant and less resilient to drought and its legacy compared to bacteria, while heat had limited overall effect on both domains. Symbiosis generated variation in resistance and resilience within the domains but did not have a universal effect; for example, notable mutualistic bacteria performed well under climate stress while mutualistic fungi did not. Our results suggest that slower-growing microbes will be more affected by climate change. Low resistance and resilience of soil fungal communities will have far-reaching consequences for ecosystems under climate change.

## Acknowledgments

We thank KSR staff throughout the years, Chris Carlson for his help in the field, and Noelle Perkins for additional soil monitoring data. This work was funded by the Government of Canada through Genome Canada and Ontario Genomics (OGI-211), NSERC Discovery grants to IE, JRS, and MEF, and the Canada Foundation for Innovation. We also acknowledge the support by the Forest and Environmental Genomics group of the Canadian Forest Service Quebec who provided white spruce seedlings and technical support during planting, and Genome Quebec for their sequencing facilities.

