## Appendix S1 for "Resistance and resilience of soil microbiomes under climate change"

Ecosphere

Climate Ecology

### Appendix S1 of ‘Resistance and resiliency of soil microbiomes under climate change’

Julia A. Boyle, Bridget K. Murphy, Ingo Ensminger, John R. Stinchcombe, Megan. E. Frederickson

*Table S1. Heat and drought exposure to the Koffler Scientific Reserve warming array plots used in this experiment. Since the plots were excavated in 2017, no prior treatments contributed. DOY= Day of the year.*

| Heat start | Heat end | Total heat duration | Drought start | Drought end | Total drought duration |
|---|---|---|---|---|---|
| August 31 2020<br>DOY=244 | December 18 2020<br>DOY=353 | 4 months/109 days | July 21 2020<br>DOY=203 | November 27 2020<br>DOY=332 | 4 months/129 days |
| August 16 2021<br>DOY=228 | January 17 2022<br>DOY=17 | 5 months/154 days | June 24 2021<br>DOY=175 | October 14 2021<br>DOY=287 | 4 months/112 days |

*Table S2. Rarefaction depth and subsequent number of amplicon sequence variants.*

| Sample | Bacteria rarefied reads (no.) | Rarefied bacterial ASVs (no.) | Fungal rarefied reads (no.) | Rarefied fungal ASVs (no.) |
|---|---|---|---|---|
| June 2021 | 134,178 | 17,836 | 109,540 | 4,250 |
| September 2021 | 129,124 | 20,198 | 127,006 | 4,064 |
| June 2022 | 284,530 | 26,588 | 180,657 | 2,194 |

*Table S3. Linear models for fungal alpha diversity indices across timepoints. 'Heat' and 'drought' estimates are relative to when heat and drought were not applied, respectively. We applied type 3 ANOVAs to the linear models to calculate the F value.*

| | | **First recovery period** | | | **Treatment** | | | **Second recovery period** | | |
|---|---|---|---|---|---|---|---|---|---|---|
| *Alpha diversity* | *Predictors* | *Estimate ±1SE* | $F_{(1,8)}$ | *p* | *Estimate ±1SE* | $F_{(1,8)}$ | *p* | *Estimate ±1SE* | $F_{(1,8)}$ | *p* |
| Shannon | Intercept | 5.09 ±0.189 | 725 | **<0.001** | 4.48 ±0.270 | 274 | **<0.001** | 4.47 ±0.260 | 295 | **<0.001** |
| | Heat | -0.082 ±0.267 | 0.095 | 0.766 | 0.485 ±0.382 | 1.61 | 0.240 | -0.245 ±0.368 | 0.878 | 0.376 |
| | Drought | -0.831 ±0.267 | 9.67 | **0.014** | -0.358 ±0.382 | 0.875 | 0.377 | -0.392 ±0.368 | 1.14 | 0.318 |
| | Heat x Drought | 1.01 ±0.378 | 7.19 | **0.028** | 0.191 ±0.541 | 0.125 | 0.733 | 0.600 ±0.520 | 1.33 | 0.282 |
| Evenness | Intercept | 0.823 ±0.028 | 862 | **<0.001** | 0.739 ±0.043 | 290 | **<0.001** | 0.673 ±0.033 | 411 | **<0.001** |
| | Heat | -7.44e-3 ±0.039 | 0.035 | 0.856 | 0.065 ±0.061 | 1.13 | 0.319 | -0.042 ±0.047 | 0.801 | 0.397 |
| | Drought | -0.131 ±0.039 | 10.9 | **0.011** | -0.060 ±0.061 | 0.958 | 0.356 | -0.037 ±0.047 | 0.627 | 0.452 |
| | Heat x Drought | 0.149 ±0.056 | 7.01 | **0.029** | 0.036 ±0.087 | 0.173 | 0.689 | 0.077 ±0.066 | 1.36 | 0.278 |
| Observed richness | Intercept | 473 ±19.9 | 562 | **<0.001** | 434 ±22.5 | 372 | **<0.001** | 768 ±56.5 | 185 | **<0.001** |
| | Heat | -19.33 ±28.2 | 0.469 | 0.513 | 46.7 ±31.8 | 2.15 | 0.181 | -79.7 ±79.8 | 0.996 | 0.348 |
| | Drought | -14.0 ±28.2 | 0.246 | 0.633 | -2.67 ±31.8 | 0.007 | 0.935 | -152 ±79.8 | 3.63 | 0.093 |
| | Heat x Drought | 55.0 ±39.9 | 1.89 | 0.206 | -7.00 ±45.0 | 0.024 | 0.880 | 104 ±113 | 0.843 | 0.385 |
| Simpson | Intercept | 0.977 ±0.009 | 111e2 | **<0.001** | 0.939 ±0.025 | 137e1 | **<0.001** | 0.957 ±0.023 | 175e1 | **<0.001** |
| | Heat | 4.51e-3 ±0.013 | 0.119 | 0.739 | 0.040 ±0.036 | 1.25 | 0.296 | -0.037 ±0.032 | 1.27 | 0.292 |
| | Drought | -0.039 ±0.013 | 8.74 | **0.018** | -0.026 ±0.036 | 0.506 | 0.497 | -0.020 ±0.032 | 0.365 | 0.562 |
| | Heat x Drought | 0.042 ±0.018 | 5.04 | 0.055 | 0.019 ±0.051 | 0.140 | 0.718 | 0.058 ±0.046 | 1.62 | 0.239 |

*Table S4. PERMANOVAs for fungal community composition across timepoints, Bray-Curtis method, 9999 permutations. 'Heat' and 'drought' estimates are relative to when heat and drought were not applied, respectively. Distance between centroids was calculated using betadisper() and dist() functions.*

| | **First recovery period** | | | **Treatment** | | | **Second recovery period** | | |
|---|---|---|---|---|---|---|---|---|---|
| *Predictors* | $F_{(1,8)}$ | *p* | *Distance between centroids* | $F_{(1,8)}$ | *p* | *Distance between centroids* | $F_{(1,8)}$ | *p* | *Distance between centroids* |
| Heat | 0.962 | 0.670 | 0.373 | 1.05 | 0.358 | 0.376 | 0.775 | 0.705 | 0.184 |
| Drought | 1.08 | 0.145 | 0.395 | 1.24 | **0.045** | 0.408 | 2.35 | **0.014** | 0.320 |
| Heat x Drought | 1.07 | 0.196 | NA | 1.06 | 0.323 | NA | 1.05 | 0.357 | NA |

*Table S5. Linear models for fungal community dynamic metrics across climate treatments. 'Heat' and 'drought' is relative to when heat and drought were not applied, respectively.*

| | **Mean rank shift** | | **Total turnover** | | **Appearances (turnover)** | | **Disappearance (turnover)** | | **Synchrony** | |
|---|---|---|---|---|---|---|---|---|---|---|
| *Predictors* | $F_{(1,19)}$ | *p* | $F_{(1,8)}$ | *p* | $F_{(1,8)}$ | *p* | $F_{(1,8)}$ | *p* | $F_{(1,8)}$ | *p* |
| Intercept | 441 | **<0.001** | 23.5 | **<0.001** | 5.90 | **0.027** | 47.9 | **<0.001** | 12.5 | **0.008** |
| Heat | 0.034 | 0.856 | 1.05 | 0.322 | 6.70 | **0.020** | 6.73 | **0.020** | 1.02 | 0.342 |
| Drought | 0.000 | 0.993 | 0.008 | 0.928 | 0.686 | 0.420 | 1.33 | 0.266 | 2.37 | 0.162 |
| Heat x Drought | 0.201 | 0.660 | 0.121 | 0.732 | 2.69 | 0.120 | 3.52 | 0.079 | 0.658 | 0.441 |
| Year pair comparison | 2.36 | 0.144 | 10.1 | **0.006** | 32.0 | **<0.001** | 24.4 | **<0.001** | | |
| Heat x Year | 0.400 | 0.536 | 1.17 | 0.296 | 7.25 | **0.016** | 7.22 | **0.016** | | |
| Drought x Year | 0.108 | 0.747 | 0.00 | 0.996 | 1.18 | 0.294 | 2.00 | 0.176 | | |
| Heat x Drought x Year | 0.178 | 0.679 | 0.160 | 0.695 | 3.07 | 0.10 | 3.93 | 0.065 | | |

*Table S6. Repeated measures ANOVAs of the relative abundance of fungal guilds. Abundance is relative to total plot reads. Mycorrhizae was log+1 transformed, while parasites and pathogens were log transformed. Endophyte and saprotroph relative abundance was already normally distributed.*

| | **Mycorrhizae abundance** | | | **Parasite abundance** | | | **Pathogen abundance** | | | **Endophyte abundance** | | | **Saprotroph abundance** | | | **Ericoid and arbuscular mycorrhizae abundance** | | | **Ectomycorrhiza abundance** | | |
|---|---|---|---|---|---|---|---|---|---|---|---|---|---|---|---|---|---|---|---|---|---|
| *Predictors* | *F* | *df* | *p* | *F* | *df* | *p* | *F* | *df* | *p* | *F* | *df* | *p* | *F* | *df* | *p* | *F* | *df* | *p* | *F* | *df* | *p* |
| Drought | 1.82 | 1,8 | 0.21 | 13.0 | 1,8 | **<0.01** | 0.140 | 1,8 | 0.718 | 1.07 | 1,8 | 0.332 | 3.21 | 1,8 | 0.111 | 1.69 | 1,8 | 0.229 | 0.853 | 1, 8 | 0.383 |
| Heat | 6.15 | 1,8 | **0.038** | 1.25 | 1,8 | 0.296 | 0.038 | 1,8 | 0.850 | 0.074 | 1,8 | 0.792 | 0.568 | 1,8 | 0.473 | 0.034 | 1,8 | 0.859 | 5.26 | 1, 8 | 0.051 |
| Drought x Heat | 3.92 | 1,8 | 0.083 | 0.757 | 1,8 | 0.409 | 1.13 | 1,8 | 0.320 | 3.12 | 1,8 | 0.116 | 0.231 | 1,8 | 0.644 | 1.63 | 1,8 | 0.238 | 2.33 | 1, 8 | 0.165 |
| Month | 13.2 | 2,22 | **<0.001** | 5.29 | 2,22 | **0.013** | 0.613 | 2,22 | 0.551 | 6.18 | 2,22 | **<0.01** | 1.25 | 2,22 | 0.306 | 2.19 | 2,22 | 0.136 | 14.5 | 2, 22 | **<0.001** |

*Table S7. Linear models for white spruce-compatible ectomycorrhiza relative abundance across timepoints. We applied type 3 ANOVAs to the models to calculate the F value.*

| | | **First recovery period** | | | **Treatment** | | | **Second recovery period** | | |
|---|---|---|---|---|---|---|---|---|---|---|
| *Response* | *Predictors* | *Estimate ±1SE* | $F_{(1,8)}$ | *p* | *Estimate ±1SE* | $F_{(1,8)}$ | *p* | *Estimate ±1SE* | $F_{(1,8)}$ | *p* |
| Relative abundance | Intercept | 6.01e-4±2.22e-4 | 7.35 | **<0.05** | 8.26e-3±1.67e-3 | 24.4 | **<0.01** | 10.9e-3±2.59e-4 | 17.6 | **<0.01** |
| | Heat | 1.78e-4±3.13e-4 | 0.324 | 0.585 | -5.96e-3±2.36e-3 | 6.36 | **0.036** | -3.59e-4±3.67e-4 | 0.955 | 0.357 |
| | Drought | -6.58e-5±3.13e-4 | 0.044 | 0.839 | -4.43e-3±2.36e-3 | 3.52 | 0.097 | 1.14e-4±3.67e-4 | 0.097 | 0.764 |
| | Heat x Drought | 2.12e-4±4.43e-4 | 0.228 | 0.646 | 5.34e-3±3.34e-3 | 2.55 | 0.149 | -4.48e-4±5.19e-4 | 0.744 | 0.413 |

*Table S8. PERMANOVAs for bacterial community composition across timepoints, Bray-Curtis method, 9999 permutations. 'Heat' and 'drought' estimates are relative to when heat and drought were not applied, respectively. Distance between centroids was calculated using betadisper() and dist() functions.*

| | **First recovery period** | | | **Treatment** | | | **Second recovery period** | | |
|---|---|---|---|---|---|---|---|---|---|
| *Predictors* | $F_{(1,8)}$ | *p* | *Distance between centroids* | $F_{(1,8)}$ | *p* | *Distance between centroids* | $F_{(1,8)}$ | *p* | *Distance between centroids* |
| Heat | 0.994 | 0.559 | 0.345 | 0.949 | 0.929 | 0.340 | 0.994 | 0.545 | 0.349 |
| Drought | 0.978 | 0.729 | 0.342 | 1.03 | 0.196 | 0.354 | 1.03 | 0.206 | 0.355 |
| Heat x Drought | 0.993 | 0.575 | NA | 1.01 | 0.365 | NA | 1.01 | 0.347 | NA |

*Table S9. Linear models for bacterial alpha diversity indices across timepoints. 'Heat' and 'drought' estimates are relative to when heat and drought were not applied, respectively. We applied type 3 ANOVAs to the linear models to calculate the F value.*

| | | **First recovery period** | | | **Treatment** | | | **Second recovery period** | | |
|---|---|---|---|---|---|---|---|---|---|---|
| *Alpha diversity* | *Predictors* | *Estimate ±1SE* | $F_{(1,8)}$ | *p* | *Estimate ±1SE* | $F_{(1,8)}$ | *p* | *Estimate ±1SE* | $F_{(1,8)}$ | *p* |
| Shannon | Intercept | 7.17 ±0.062 | 133e2 | **<0.001** | 7.35 ±0.146 | 254e1 | **<0.001** | 7.55 ±0.021 | 1.32e5 | **<0.001** |
| | Heat | 0.080 ±0.088 | 0.831 | 0.389 | -0.089 ±0.206 | 0.190 | 0.674 | 0.051 ±0.029 | 3.02 | 0.120 |
| | Drought | 0.093 ±0.088 | 1.11 | 0.322 | -0.091 ±0.206 | 0.196 | 0.670 | 0.089 ±0.029 | 9.30 | **0.016** |
| | Heat x Drought | -0.172 ±0.125 | 1.91 | 0.204 | 0.135 ±0.292 | 0.215 | 0.656 | -0.073 ±0.041 | 3.12 | 0.115 |
| Evenness | Intercept | 0.933 ±0.001 | 3.47e5 | **<0.001** | 0.934 ±0.002 | 1.49e5 | **<0.001** | 0.927 ±0.001 | 4.45e5 | **<0.001** |
| | Heat | 7.29e-4 ±0.002 | 0.106 | 0.753 | -1.16e-3 ±0.003 | 0.115 | 0.743 | 2.11e-3 ±0.002 | 1.16 | 0.313 |
| | Drought | 4.27e-3 ±0.002 | 3.64 | 0.093 | 7.16e-4 ±0.003 | 0.044 | 0.839 | 4.40e-4 ±0.002 | 5.01e-2 | 0.829 |
| | Heat x Drought | -3.64e-3 ±0.003 | 1.32 | 0.284 | 1.97e-3 ±0.005 | 0.166 | 0.695 | -1.80e-3 ±0.003 | 4.21e-1 | 0.535 |
| Observed richness | Intercept | 220e1 ±144 | 234 | **<0.001** | 265e1 ±346 | 58.6 | **<0.001** | 348e1 ±109 | 102e1 | **<0.001** |
| | Heat | 182 ±203 | 0.801 | 0.397 | -221 ±489 | 0.203 | 0.664 | 128 ±154 | 0.689 | 0.431 |
| | Drought | 138 ±203 | 0.460 | 0.517 | -253 ±489 | 0.268 | 0.619 | 349 ±154 | 5.11 | 0.054 |
| | Heat x Drought | -354 ±287 | 1.52 | 0.253 | 369 ±692 | 0.284 | 0.608 | -240 ±218 | 1.21 | 0.303 |
| Simpson | Intercept | 0.999 ±8.03e-5 | 1.55e8 | **<0.001** | 0.999 ±1.78e-4 | 3.13e7 | **<0.001** | 0.999 ±2.84e-5 | 1.24e9 | **<0.001** |
| | Heat | 4.99e-5 ±1.14e-4 | 0.194 | 0.672 | -9.34e-5 ±2.53e-4 | 0.137 | 0.721 | 2.59e-5 ±4.01e-5 | 0.417 | 0.537 |
| | Drought | 1.61e-4 ±1.14e-4 | 2.02 | 0.193 | -6.91e-5 ±2.53e-4 | 7.50e-2 | 0.791 | 6.93e-5 ±4.01e-5 | 2.98 | 0.123 |
| | Heat x Drought | -1.97e-4 ±1.61e-4 | 1.51 | 0.254 | 1.21e-4 ±3.57e-4 | 0.114 | 0.744 | -4.44e-5 ±5.68e-5 | 0.611 | 0.457 |

*Table S10. Linear models for bacterial community dynamic metrics across climate treatments. 'Heat' and 'drought' estimates are relative to when heat and drought were not applied, respectively.*

| | **Mean rank shift** | | **Total turnover** | | **Appearances (turnover)** | | **Disappearance (turnover)** | | **Synchrony** | |
|---|---|---|---|---|---|---|---|---|---|---|
| *Predictors* | $F_{(1,19)}$ | $p$ | $F_{(1,8)}$ | $p$ | $F_{(1,8)}$ | $p$ | $F_{(1,8)}$ | $p$ | $F_{(1,8)}$ | $p$ |
| Intercept | 121 | **<0.001** | 24.1 | **<0.001** | 4.94 | **0.041** | 1.31 | 0.269 | 531e1 | **<0.001** |
| Heat | 0.001 | 0.982 | 0.250 | 0.62 | 0.354 | 0.560 | 0.047 | 0.831 | 0.458 | 0.518 |
| Drought | 0.023 | 0.881 | 0.795 | 0.386 | 2.18 | 0.159 | 0.581 | 0.457 | 0.896 | 0.372 |
| Heat x Drought | 0.004 | 0.950 | 0.256 | 0.620 | 2.81 | 0.113 | 1.42 | 0.252 | 0.009 | 0.925 |
| Year pair comparison | 1.09 | 0.311 | 1.51 | 0.237 | 0.647 | 0.433 | 0.004 | 0.953 | | |
| Heat x Year | 0.589 | 0.454 | 0.627 | 0.440 | 0.579 | 0.458 | 0.032 | 0.860 | | |
| Drought x Year | 0.371 | 0.551 | 1.53 | 0.234 | 2.62 | 0.125 | 0.447 | 0.513 | | |
| Heat x Drought x Year | 0.144 | 0.709 | 0.565 | 0.463 | 3.16 | 0.094 | 1.26 | 0.277 | | |

*Table S11. Phylogenetic signal in the differential abundance of genera, using Pagel's λ. 'Heat' and 'drought' estimates are relative to when heat and drought were not applied, respectively. Loglikelihood ratio test is compared to when λ=0.*

| | **First recovery period** | | | | **Active treatment** | | | | **Second recovery period** | | | |
|---|---|---|---|---|---|---|---|---|---|---|---|---|
| | λ | *Loglikelihood ratio* | *Loglikelihood of λ* | *p* | λ | *Loglikelihood ratio* | *Loglikelihood of λ* | *p* | λ | *Loglikelihood ratio* | *Loglikelihood of λ* | *p* |
| Heat | 7.33 e-5 | -0.004 | -258 | 1.00 | 0.083 | 2.13 | -276 | 0.144 | 0.308 | 17.9 | -253 | **<0.001** |
| Drought | 7.33 e-5 | -0.003 | -252 | 1.00 | 0.022 | 0.327 | -276 | 0.568 | 0.190 | 7.71 | -240 | **<0.01** |

*Table S12. Linear models for phytopathogenic bacteria alpha diversity indices across timepoints. We applied type 3 ANOVAs to the linear models to calculate the F value.*

| | | **First recovery period** | | | **Treatment** | | | **Second recovery period** | | |
|---|---|---|---|---|---|---|---|---|---|---|
| *Alpha diversity* | *Predictors* | *Estimate ±1SE* | $F_{(1,8)}$ | *p* | *Estimate ±1SE* | $F_{(1,8)}$ | *p* | *Estimate ±1SE* | $F_{(1,8)}$ | *p* |
| Shannon | Intercept | 3.12 ±0.117 | 710 | **<0.001** | 1.16 ±0.072 | 265 | **<0.001** | 3.42 ±0.079 | 187e1 | **<0.001** |
| | Heat | 0.037 ±0.166 | 0.050 | 0.829 | 0.006 ±0.101 | 0.004 | 0.953 | 0.028 ±0.112 | 0.064 | 0.807 |
| | Drought | 0.085 ±0.166 | 0.266 | 0.620 | 0.014 ±0.101 | 0.019 | 0.895 | 0.202 ±0.112 | 3.26 | 0.109 |
| | Heat x Drought | -0.14 ±0.234 | 0.356 | 0.567 | 0.056 ±0.143 | 0.156 | 0.704 | -0.109 ±0.158 | 0.473 | 0.511 |
| Evenness | Intercept | 0.889 ±0.017 | 259e1 | **<0.001** | 0.886 ±0.021 | 26.6 | **<0.001** | 0.868 ±0.005 | 257e2 | **<0.001** |
| | Heat | -0.021 ±0.025 | 0.740 | 0.415 | -0.003 ±0.029 | 0.006 | 0.941 | -0.026 ±0.008 | 11.8 | **0.009** |
| | Drought | -0.015 ±0.025 | 0.358 | 0.566 | -0.015 ±0.029 | 0.247 | 0.633 | 0.010 ±0.008 | 1.64 | 0.237 |
| | Heat x Drought | 0.025 ±0.035 | 0.521 | 0.491 | 0.039 ±0.041 | 0.879 | 0.376 | 0.015 ±0.011 | 1.99 | 0.196 |
| Observed richness | Intercept | 34.0 ±3.26 | 109 | **<0.001** | 40.3 ±6.53 | 427 | **<0.001** | 51.7 ±4.32 | 143 | **<0.001** |
| | Heat | 4.33 ±4.61 | 0.883 | 0.375 | -0.333 ±9.23 | 0.007 | 0.935 | 9.33 ±6.11 | 2.34 | 0.165 |
| | Drought | 5.33 ±4.61 | 1.34 | 0.281 | 3.00 ±9.23 | 0.167 | 0.694 | 10.67 ±6.11 | 3.05 | 0.119 |
| | Heat x Drought | -8.67 ±6.52 | 1.77 | 0.221 | 2.33 ±13.1 | 0.020 | 0.892 | -11.7 ±8.64 | 1.83 | 0.214 |
| Simpson | Intercept | 0.938 ±0.011 | 671e1 | **<0.001** | 0.929 ±0.021 | 10.4 | **0.012** | 0.945 ±0.006 | 285e2 | **<0.001** |
| | Heat | -6.50e-3 ±0.016 | 0.161 | 0.700 | 0.007 ±0.030 | 0.061 | 0.811 | -0.010 ±0.008 | 1.76 | 0.221 |
| | Drought | -3.61e-3 ±0.016 | 0.050 | 0.829 | -0.001 ±0.030 | 0.001 | 0.979 | 0.010 ±0.008 | 1.84 | 0.212 |
| | Heat x Drought | -1.65e-4 ±0.023 | 0.00 | 0.994 | 0.020 ±0.042 | 0.216 | 0.654 | 0.004 ±0.011 | 0.134 | 0.724 |

*Table S13. PERMANOVAs for phytopathogen community composition across timepoints, Bray-Curtis method, 9999 permutations. 'Heat' and 'drought' estimates are relative to when heat and drought were not applied, respectively. Betadisper and dist functions used to calculate distance between centroids.*

| | **First recovery period** | | | **Treatment** | | | **Second recovery period** | | |
|---|---|---|---|---|---|---|---|---|---|
| *Predictors* | $F_{(1,8)}$ | *p* | *Distance between centroids* | $F_{(1,8)}$ | *p* | *Distance between centroids* | $F_{(1,8)}$ | *p* | *Distance between centroids* |
| Heat | 0.947 | 0.534 | 0.320 | 0.790 | 0.846 | 0.319 | 0.975 | 0.487 | 0.328 |
| Drought | 0.918 | 0.599 | 0.315 | 0.799 | 0.823 | 0.321 | 1.75 | **0.019** | 0.439 |
| Heat x Drought | 1.09 | 0.319 | NA | 0.979 | 0.501 | NA | 1.13 | 0.229 | NA |

*Table S14. Linear models for plant beneficial bacteria alpha diversity indices across timepoints. We applied type 3 ANOVAs to the linear models to calculate the F value.*

| | | **First recovery period** | | | **Treatment** | | | **Second recovery period** | | |
|---|---|---|---|---|---|---|---|---|---|---|
| *Alpha diversity* | *Predictors* | *Estimate ±1SE* | $F_{(1,8)}$ | *p* | *Estimate ±1SE* | $F_{(1,8)}$ | *p* | *Estimate ±1SE* | $F_{(1,8)}$ | *p* |
| Shannon | Intercept | 4.70 ±0.046 | 102e2 | **<0.001** | 1.57 ±0.029 | 296e1 | **<0.001** | 5.07 ±0.036 | 201e2 | **<0.001** |
| | Heat | 0.030 ±0.066 | 0.213 | 0.656 | 0.005 ±0.041 | 0.018 | 0.897 | 0.050 ±0.051 | 0.962 | 0.356 |
| | Drought | 0.116 ±0.066 | 3.09 | 0.117 | -0.007 ±0.041 | 0.029 | 0.870 | 0.172 ±0.051 | 11.6 | **0.009** |
| | Heat x Drought | -0.149 ±0.093 | 2.56 | 0.148 | 0.023 ±0.058 | 0.154 | 0.705 | -0.112 ±0.072 | 2.48 | 0.154 |
| Evenness | Intercept | 0.917 ±0.006 | 271e2 | **<0.001** | 0.926 ±0.004 | 700e2 | **<0.001** | 0.915 ±0.002 | 1.36e5 | **<0.001** |
| | Heat | -0.004 ±0.008 | 0.212 | 0.658 | -0.005 ±0.005 | 1.05 | 0.336 | -0.004 ±0.004 | 1.59 | 0.243 |
| | Drought | 0.001 ±0.008 | 0.015 | 0.907 | -0.003 ±0.005 | 0.286 | 0.607 | 0.003 ±0.004 | 0.554 | 0.478 |
| | Heat x Drought | 0.006 ±0.011 | 0.253 | 0.629 | 0.007 ±0.007 | 0.954 | 0.357 | 0.004 ±0.005 | 0.786 | 0.401 |
| Observed richness | Intercept | 170 ±10.2 | 279 | **<0.001** | 186 ±26.4 | 49.3 | **<0.001** | 254 ±6.86 | 137e1 | **<0.001** |
| | Heat | 8.33 ±14.4 | 0.336 | 0.578 | 11.0 ±37.4 | 0.087 | 0.776 | 21.3 ±9.70 | 4.84 | 0.059 |
| | Drought | 20.7 ±14.4 | 2.07 | 0.188 | -6.00 ±37.4 | 0.026 | 0.877 | 47.3 ±9.70 | 23.8 | **0.001** |
| | Heat x Drought | -33.0 ±20.3 | 2.64 | 0.143 | 19.0 ±52.9 | 0.129 | 0.729 | -41.0 ±13.7 | 8.94 | **0.017** |
| Simpson | Intercept | 0.987 ±0.001 | 2.12e6 | **<0.001** | 0.988 ±0.002 | 2.99e5 | **<0.001** | 0.99 ±3.74e-4 | 7.02e6 | **<0.001** |
| | Heat | -3.10e-4 | 0.104 | 0.755 | 5.33e-4 ±0.003 | 0.044 | 0.840 | -2.48e-4±5.29e-4 | 0.220 | 0.651 |
| | Drought | 9.06e-4 | 0.892 | 0.372 | -2.29e-4 ±0.003 | 0.008 | 0.931 | 0.002 ±5.29e-4 | 12.3 | **0.008** |
| | Heat x Drought | -9.28e-4 | 0.468 | 0.513 | 0.001 ±0.004 | 0.115 | 0.744 | -4.51e-4 ±7.48e-4 | 0.365 | 0.563 |

*Table S15. PERMANOVAs for plant beneficial bacteria community composition across timepoints, Bray-Curtis method, 9999 permutations. 'Heat' and 'drought' estimates are relative to when heat and drought were not applied, respectively. Betadisper and dist functions used to calculate distance between centroids.*

| | **First recovery period** | | | **Treatment** | | | **Second recovery period** | | |
|---|---|---|---|---|---|---|---|---|---|
| *Predictors* | $F_{(1,8)}$ | *p* | *Distance between centroids* | $F_{(1,8)}$ | *p* | *Distance between centroids* | $F_{(1,8)}$ | *p* | *Distance between centroids* |
| Heat | 1.13 | 0.104 | 0.363 | 0.937 | 0.772 | 0.340 | 0.897 | 0.848 | 0.329 |
| Drought | 0.911 | 0.828 | 0.327 | 1.10 | 0.127 | 0.369 | 1.02 | 0.388 | 0.352 |
| Heat x Drought | 0.995 | 0.514 | NA | 0.918 | 0.848 | NA | 1.06 | 0.253 | NA |

*Table S16. Linear models for rhizobia alpha diversity indices across timepoints. 'Heat' and 'drought' estimates are relative to when heat and drought were not applied, respectively. We applied type 3 ANOVAs to the linear models to calculate the F value.*

| | | **First recovery period** | | | **Treatment** | | | **Second recovery period** | | |
|---|---|---|---|---|---|---|---|---|---|---|
| *Alpha diversity* | *Predictors* | *Estimate ±1SE* | $F_{(1,8)}$ | *p* | *Estimate ±1SE* | $F_{(1,8)}$ | *p* | *Estimate ±1SE* | $F_{(1,8)}$ | *p* |
| Shannon | Intercept | 2.54 ±0.085 | 902 | **<0.001** | 2.51 ±0.095 | 696 | **<0.001** | 2.48 ±0.138 | 322 | **<0.001** |
| | Heat | -0.055 ±0.119 | 0.212 | 0.657 | 0.421 ±0.135 | 9.78 | **0.014** | 0.165 ±0.196 | 0.71 | 0.425 |
| | Drought | 0.227 ±0.119 | 3.59 | 0.095 | 0.310 ±0.135 | 5.29 | **0.050** | 0.477 ±0.196 | 5.94 | **0.041** |
| | Heat x Drought | 0.055 ±0.169 | 0.107 | 0.752 | -0.319 ±0.191 | 2.81 | 0.132 | -0.303 ±0.277 | 1.19 | 0.306 |
| Evenness | Intercept | 0.863 ±0.017 | 244e1 | **<0.001** | 0.889 ±0.012 | 567e1 | **<0.001** | 0.821 ±0.022 | 138e1 | **<0.001** |
| | Heat | -0.021 ±0.025 | 0.751 | 0.411 | 0.023 ±0.017 | 1.85 | 0.211 | 0.017 ±0.031 | 0.306 | 0.595 |
| | Drought | 0.029 ±0.025 | 1.44 | 0.265 | 0.019 ±0.017 | 1.26 | 0.294 | 0.071 ±0.031 | 5.22 | 0.052 |
| | Heat x Drought | 0.026 ±0.035 | 0.535 | 0.486 | -0.022 ±0.023 | 0.908 | 0.368 | -0.046 ±0.044 | 1.09 | 0.326 |
| Observed | Intercept | 19.3 ±1.88 | 106 | **<0.001** | 17.3 ±2.21 | 61.8 | **<0.001** | 20.7 ±2.30 | 80.9 | **<0.001** |
| richness | Heat | -1.92e-16 ±2.65 | 0.00 | 1.00 | 8.00 ±3.12 | 6.58 | **0.033** | 3.00 ±3.25 | 0.853 | 0.383 |
| | Drought | 3.00 ±2.66 | 1.28 | 0.291 | 5.33 ±3.12 | 2.93 | 0.126 | 7.00 ±3.25 | 4.64 | 0.063 |
| | Heat x Drought | -7.18e-15 ±3.76 | 0.00 | 1.00 | -5.33 ±4.41 | 1.46 | 0.261 | -4.00 ±4.59 | 0.758 | 0.409 |
| Simpson | Intercept | 0.891 ±0.014 | 398e1 | **<0.001** | 0.897 ±0.007 | 150e2 | **<0.001** | 0.878 ±0.017 | 275e1 | **<0.001** |
| | Heat | -0.017 ±0.020 | 0.691 | 0.429 | 0.036 ±0.010 | 12.1 | **0.008** | 0.012 ±0.024 | 0.271 | 0.617 |
| | Drought | 0.025 ±0.020 | 1.62 | 0.239 | 0.031 ±0.010 | 8.86 | **0.018** | 0.055 ±0.024 | 5.46 | **0.047** |
| | Heat x Drought | 0.015 ±0.028 | 0.283 | 0.609 | -0.029 ±0.015 | 4.16 | 0.075 | -0.032 ±0.034 | 0.893 | 0.372 |

*Table S17. PERMANOVAs for rhizobia community composition across timepoints, Bray-Curtis method, 9999 permutations. 'Heat' and 'drought' estimates are relative to when heat and drought were not applied, respectively. Betadisper and dist functions used to calculate distance between centroids.*

| | **First recovery period** | | | **Treatment** | | | **Second recovery period** | | |
|---|---|---|---|---|---|---|---|---|---|
| *Predictors* | $F_{(1,8)}$ | *p* | *Distance between centroids* | $F_{(1,8)}$ | *p* | *Distance between centroids* | $F_{(1,8)}$ | *p* | *Distance between centroids* |
| Heat | 1.21 | 0.235 | 0.351 | 0.997 | 0.431 | 0.330 | 0.872 | 0.681 | 0.326 |
| Drought | 0.935 | 0.522 | 0.309 | 1.19 | 0.218 | 0.362 | 0.894 | 0.638 | 0.330 |
| Heat x Drought | 1.78 | **0.024** | NA | 1.08 | 0.336 | NA | 1.47 | 0.063 | NA |

*Table S18. Network analysis summary table. Genus level analysis of combined bacteria and fungal datasets. SparCC correlation cutoff of 0.8. The number of significant correlations was determined by permuting the correlations 100 times and adjusting for false discovery rate.*

| *Network property* | **Active treatment** | | | |
|---|---|---|---|---|
| | *Control* | *Heat* | *Drought* | *Heat + Drought* |
| Node number | 331 | 367 | 352 | 361 |
| Edge number | 23516 | 28529 | 26666 | 27560 |
| Walktrap cluster number | 3 | 3 | 3 | 3 |
| Global clustering coefficient | 0.756 | 0.755 | 0.779 | 0.764 |
| Modularity | 0.350 | 0.372 | 0.369 | 0.406 |
| Top 5 'hub' genera | 1.*Emericellopsis*<br>2.*Pirellula*<br>3.*Gemmata*<br>4.*Chthoniobacter*<br>5.*DA101* | 1.*Lecythophora*<br>2.*Dactylonectria*<br>3.*Rhodoplanes*<br>4.*Hyphomicrobium*<br>5.*Mesorhizobium* | 1.*Plectosphaerella*<br>2.*Bacillus*<br>3.*Candidatus Nitrososphaera*<br>4.*Polytolypa*<br>5.*Phenylobacterium* | 1.*Gemmata*<br>2.*Demequina*<br>3.*Sterigmatobotrys*<br>4.*Thelonectria*<br>5.*Balneimonas* |
| No. of sig. bacteria-bacteria correlations | 6<br>(16% positive) | 6<br>(50% positive) | 0 | 0 |
| No. of sig. bacteria-fungal correlations | 22<br>(18% positive) | 17<br>(47% positive) | 1<br>(100% positive) | 5<br>(40% positive) |
| No. of sig. fungi-fungi correlations | 9<br>(33% positive) | 20<br>(50% positive) | 0 | 4<br>(0% positive) |

*Table S19. Hub score in control conditions is linked to the log fold response of genera to heat and drought. Below are results from the MANOVA overall as well as the significance of individual response factors using summary.aov().*

| *Predictor* | *Overall model* | | | *Response to drought* | | *Response to heat* | |
|---|---|---|---|---|---|---|---|
| | *Pillai* | $F_{(1,59)}$ | $p$ | $F_{(1,59)}$ | $p$ | $F_{(1,59)}$ | $p$ |
| *Hub score* | 0.151 | 5.16 | **0.009** | 9.19 | **0.004** | 3.31 | 0.074 |
