## Appendix S2 for "Resistance and resilience of soil microbiomes under climate change"

Ecosphere

Climate Ecology

### Appendix S2 of ‘Resistance and resiliency of soil microbiomes under climate change’

Julia A. Boyle, Bridget K. Murphy, Ingo Ensminger, John R. Stinchcombe, Megan. E. Frederickson

**Methods and Supplementary for Soil Sensor Data for KSR 2020 and 2021**

Each plot was equipped with a soil sensor (Models 5TM and TE11, METER Group, Pullman, WA, USA) in the center at a depth of 15-20cm to record soil volumetric water content (VWC) and soil temperature every 30 minutes using a CR1000 datalogger (Campbell Scientific Inc., Edmonton, AB, Canada). Sensors and dataloggers were repaired or replaced as needed throughout the experimental time frames in 2020 and 2021, however, there are periods where equipment malfunctions resulted in partial losses in soil sensor data (Table 1). The heaters still ran through these partial data losses.

Several manual soil VWC measurements were taken during the experiments in 2020 and 2021 using a HydroSense system with the CS620 sensor (Campbell Scientific Inc., Edmonton, AB, Canada) at a depth of 20cm to match the depth of the automatic soil sensors, with seven manual soil VWC measurement time points in 2020 and eight time points in 2021. At each time point, we measured soil VWC at seven points within each plot and calculated a mean value for each plot (Figure 1).

For each year, a correction factor was calculated per plot to correct for sensor drift and aging of the 5TM and TE11 probes that were permanently installed in the experimental plots. The correction factors were calculated as the ratio between the manual soil VWC means and the single automatic soil VWC sensor reading at multiple time points per experiment (Figures 2 and 3). The automatic VWC sensor readings were then multiplied by the calculated correction factor across the full sensor datasets. Both uncorrected and corrected daily soil volumetric content (VWC) values per treatment are shown in Figures 2 and 3.

**Table S1. Record of soil sensor or datalogger issues and repairs during the field experiment.**

| Field Seasons | Dates | Datalogger or Sensor Issue (Description) |
|---|---|---|
| 2020 | July 20-September 5 | Plot 10 - Malfunctioning soil sensor, no soil moisture and soil temperature recording. |
| | July 20-November 21 | Plot 7 – Malfunctioning soil sensor, no soil temperature recording. |
| | October 13-16 | Datalogger malfunctioning, no soil moisture and soil temperature recording. |
| | December 5-15 | Datalogger malfunctioning, no soil moisture and soil temperature recording for plots 7-12. |

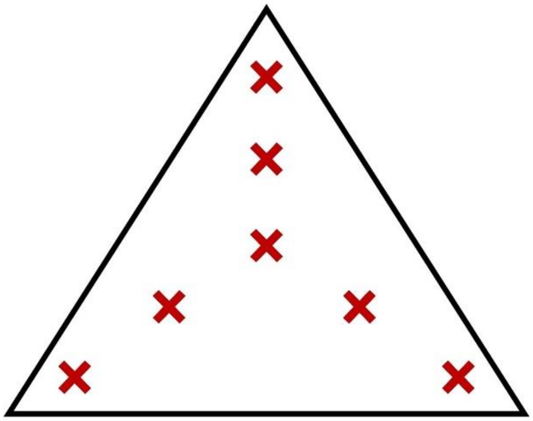

**Figure S1**. **Approximate points where manual soil volumetric water content (VWC) was measured and then averaged from in each plot.**

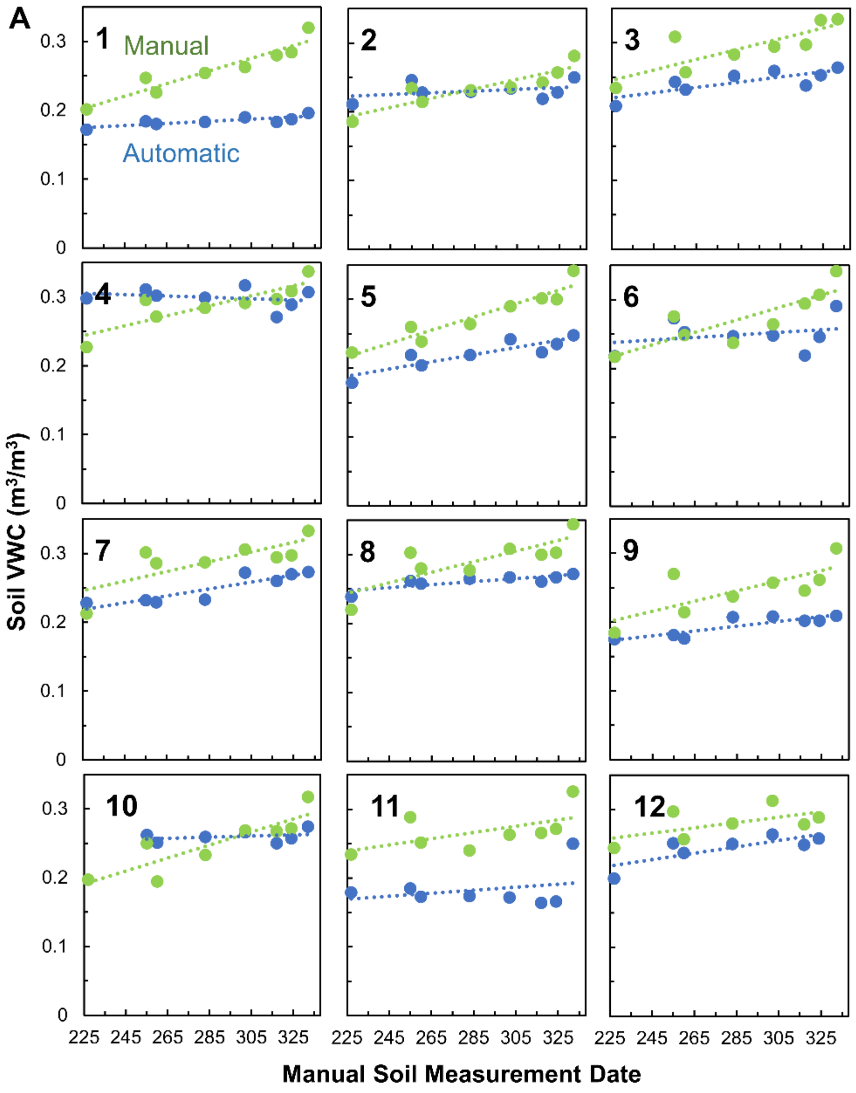


**KSR 2020**

**B**

| Plot | Corr. Factor |
|---|---|
| 1 | 1.407 |
| 2 | 1.021 |
| 3 | 1.201 |
| 4 | 0.966 |
| 5 | 1.254 |
| 6 | 1.096 |
| 7 | 1.160 |
| 8 | 1.117 |
| 9 | 1.267 |
| 10 | 0.961 |
| 11 | 1.463 |
| 12 | 1.165 |

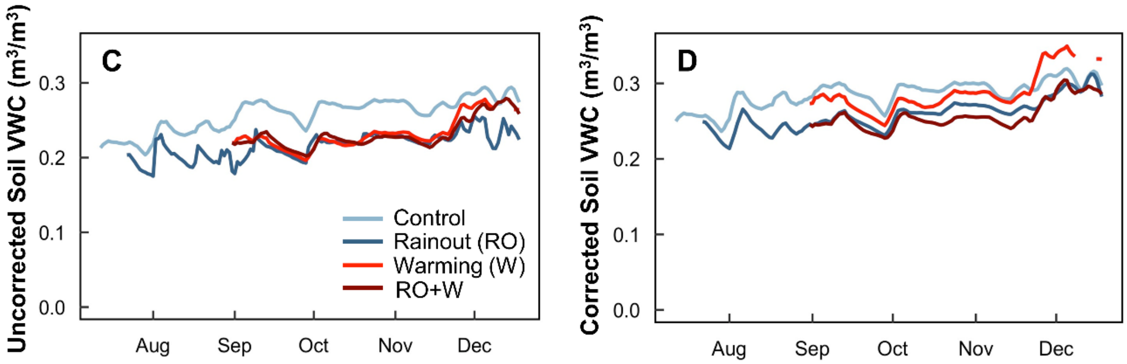


**Figure S2. Automatic soil sensor corrections applied for the 2020 experimental time frame from July 21 to Dec 18 at the Koffler Scientific Reserve in Ontario, Canada. (A)** Comparison of the mean manual soil volumetric water content (VWC; green) against the automatic soil sensor (blue) at the same time and date across the experiment within each plot. Plot numbers can be found in the upper left corners of the subplots. **(B)** The calculated correction factors were determined by the ratio of the mean manual soil VWC to the automatic soil VWC points across all time points per plot. **(C)** the 5-day running average of daily soil VWC across each treatment (blue, control; dark blue, rainout; light red, warming; dark red, warming combined with rainout) across the experimental time frame in 2020 before the correction factors were applied. **(D)** the 5-day running average of the corrected daily automatic soil VWC across each treatment across the experimental time frame in 2020 using the correction factors found in Part **B** of the figure.

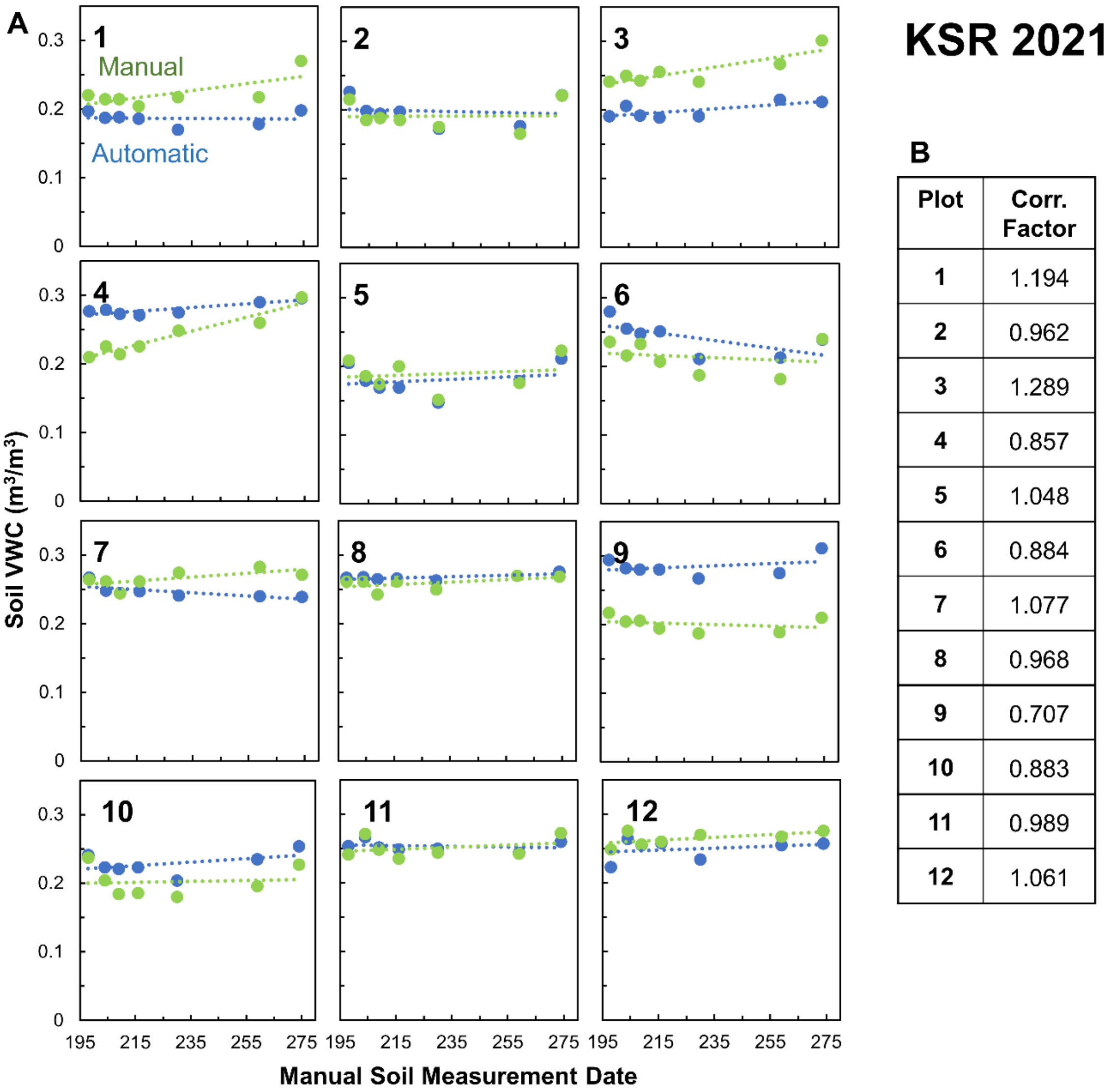


**B**

| Plot | Corr. Factor |
|---|---|
| 1 | 1.194 |
| 2 | 0.962 |
| 3 | 1.289 |
| 4 | 0.857 |
| 5 | 1.048 |
| 6 | 0.884 |
| 7 | 1.077 |
| 8 | 0.968 |
| 9 | 0.707 |
| 10 | 0.883 |
| 11 | 0.989 |
| 12 | 1.061 |

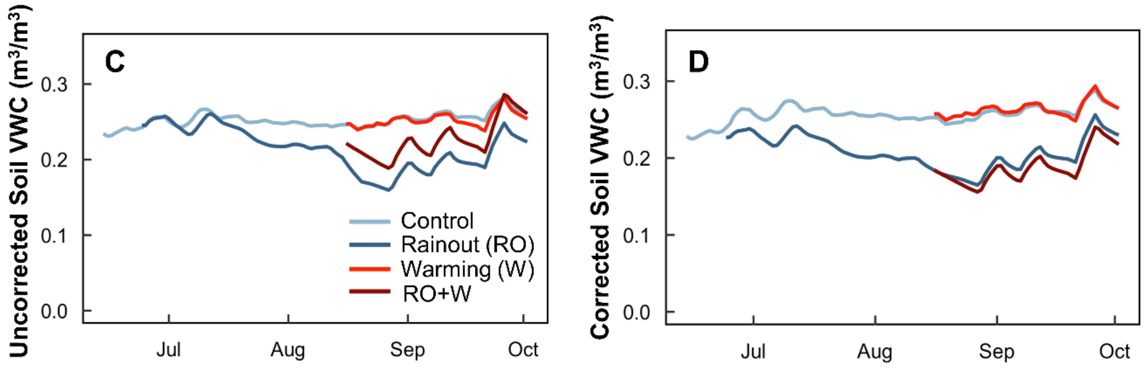


**Figure S3. Automatic soil sensor corrections applied for the 2021 experimental time frame from June 24 to October 2 at the Koffler Scientific Reserve in Ontario, Canada. (A)** Comparison of the mean manual soil volumetric water content (VWC; green) against the automatic soil sensor (blue) at the same time and date across the experiment within each plot. Plot numbers can be found in the upper left corners of the subplots. **(B)** The calculated correction factors were determined by the ratio of the mean manual soil VWC to the automatic soil VWC points across all time points per plot. **(C)** the 5-day running average of daily soil VWC across each treatment (blue, control; dark blue, rainout; light red, warming; dark red, warming combined with rainout) across the experimental time frame in 2020 before the correction factors were applied. **(D)** the 5-day running average of the corrected daily automatic soil VWC across each treatment across the experimental time frame in 2020 using the correction factors found in Part **B** of the figure.

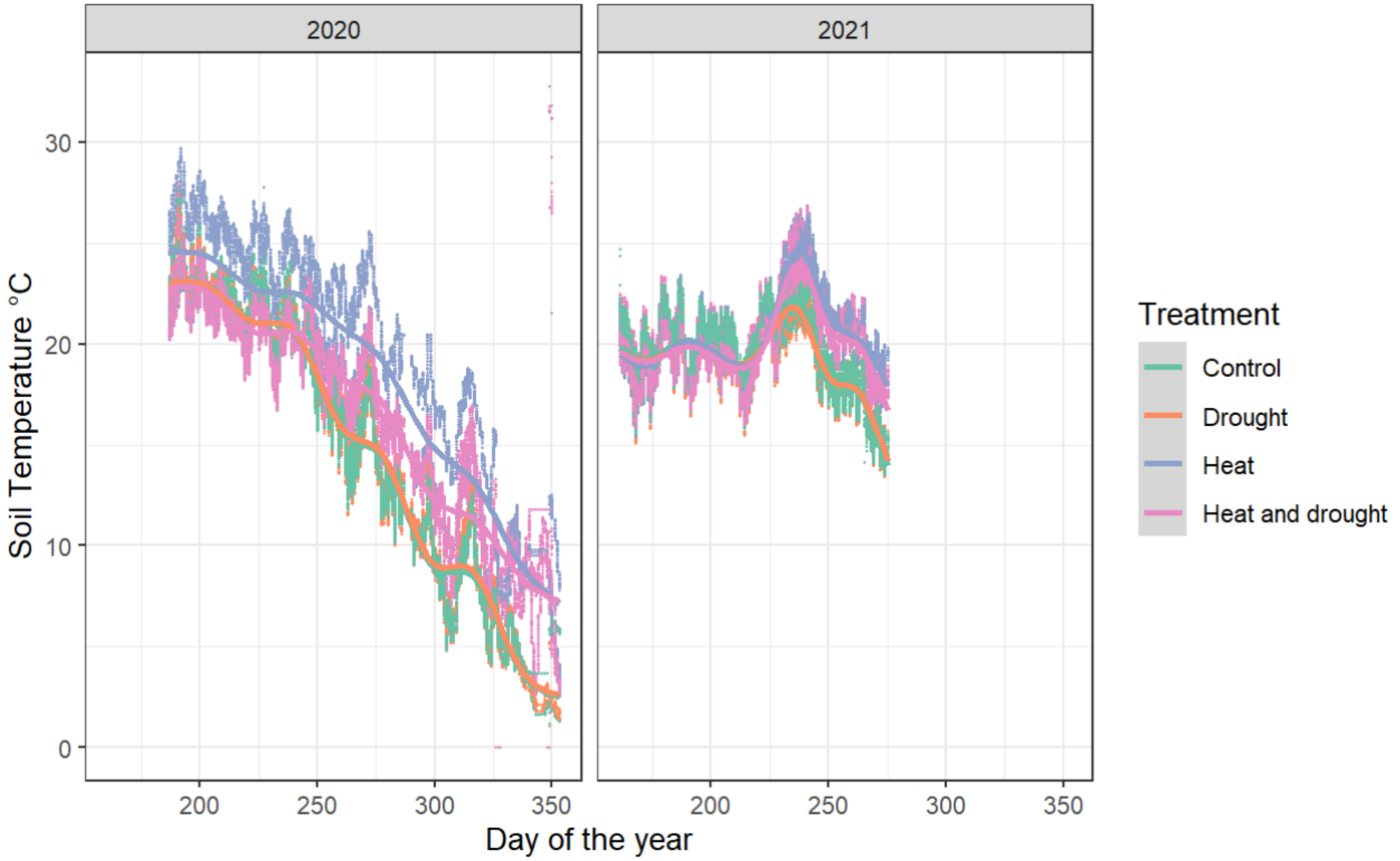


**Figure S4. Soil temperature (°C) in plots grouped by climate treatment in 2020 and 2021.** We recorded soil temperature every 30 minutes using a CR1000 datalogger during active treatment. We sampled soil in recovery periods on day 163 in 2020 and 156 in 2021, and during active treatment on day 258 in 2020.
