## Appendix S3 for "Resistance and resilience of soil microbiomes under climate change"

Ecosphere

Climate Ecology

### Appendix S3 of ‘Resistance and resiliency of soil microbiomes under climate change’

Julia A. Boyle, Bridget K. Murphy, Ingo Ensminger, John R. Stinchcombe, Megan. E. Frederickson

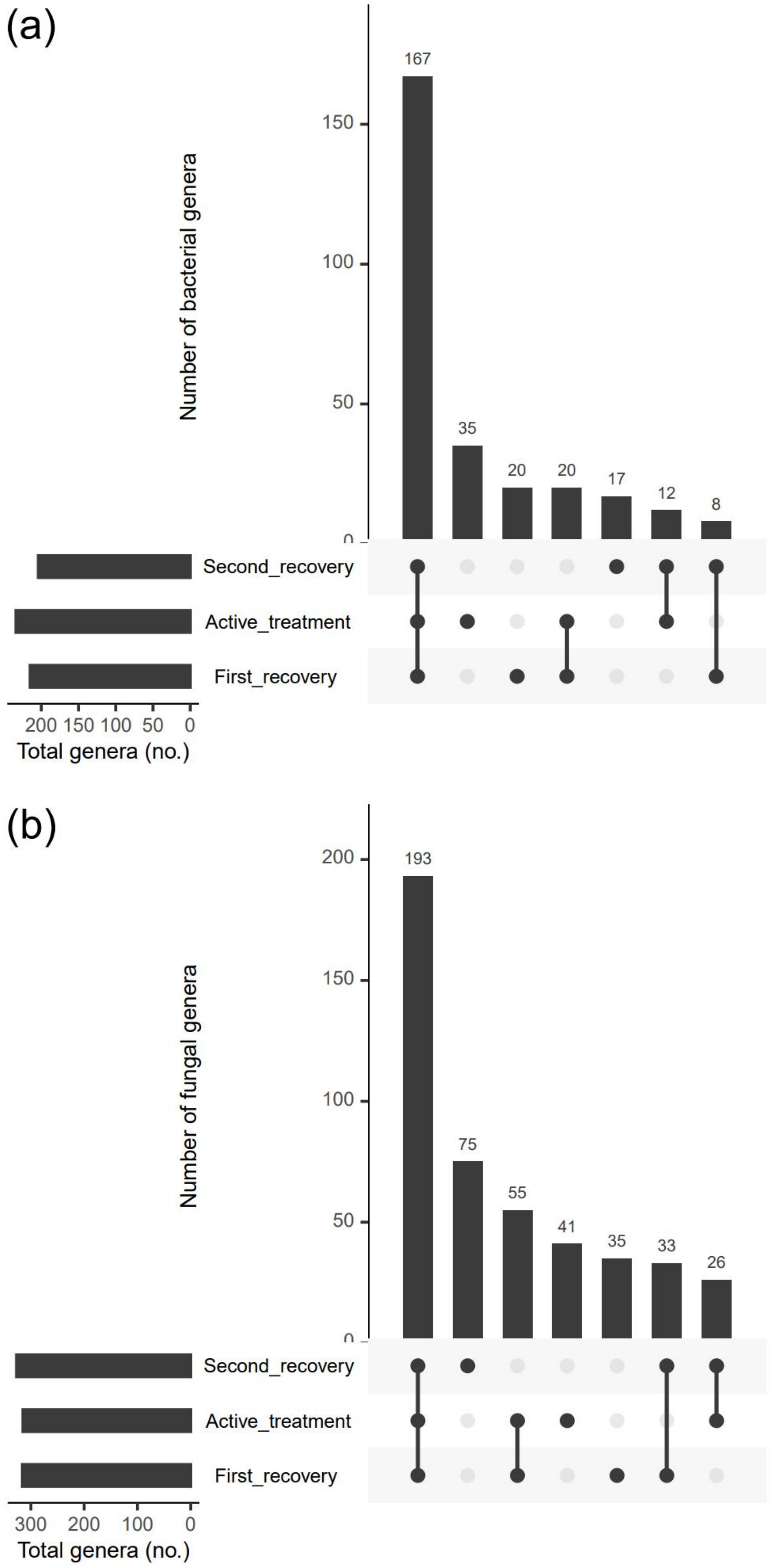


Figure S1. Overlap in bacterial (a) and fungal (b) genera across active treatment and recovery periods.

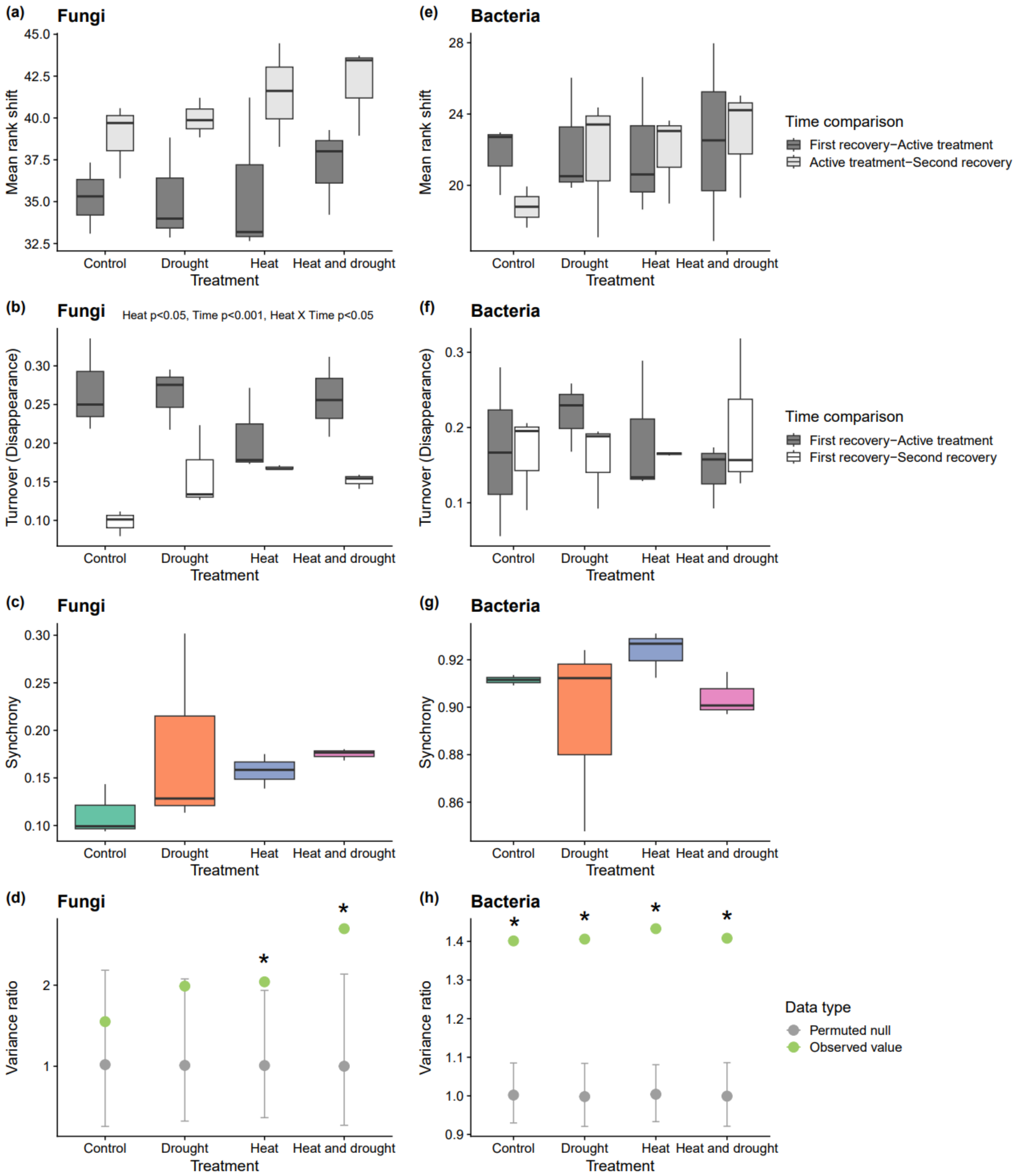


Figure S2. Community dynamics of bacterial and fungal communities across climate treatments and time. Mean rank shift, turnover and synchrony are shown using boxplots. Variance ratio plots show null permuted values with 95% confidence intervals. Only significant main effects of heat, drought, time, and their interactions are displayed on the panels. Note the scale differences between pairs of plots.

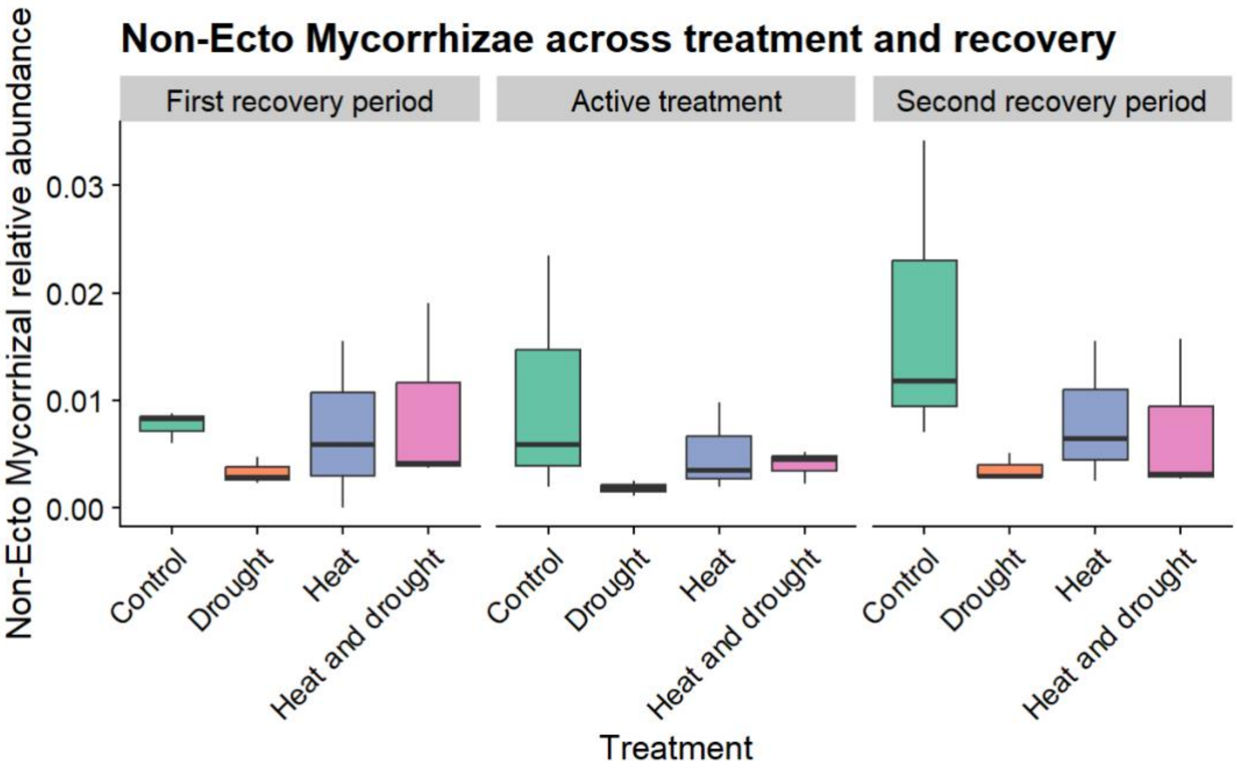

Figure S3. Relative abundance of non-ectomycorrhizal mycorrhizal fungi across climate treatments and time.

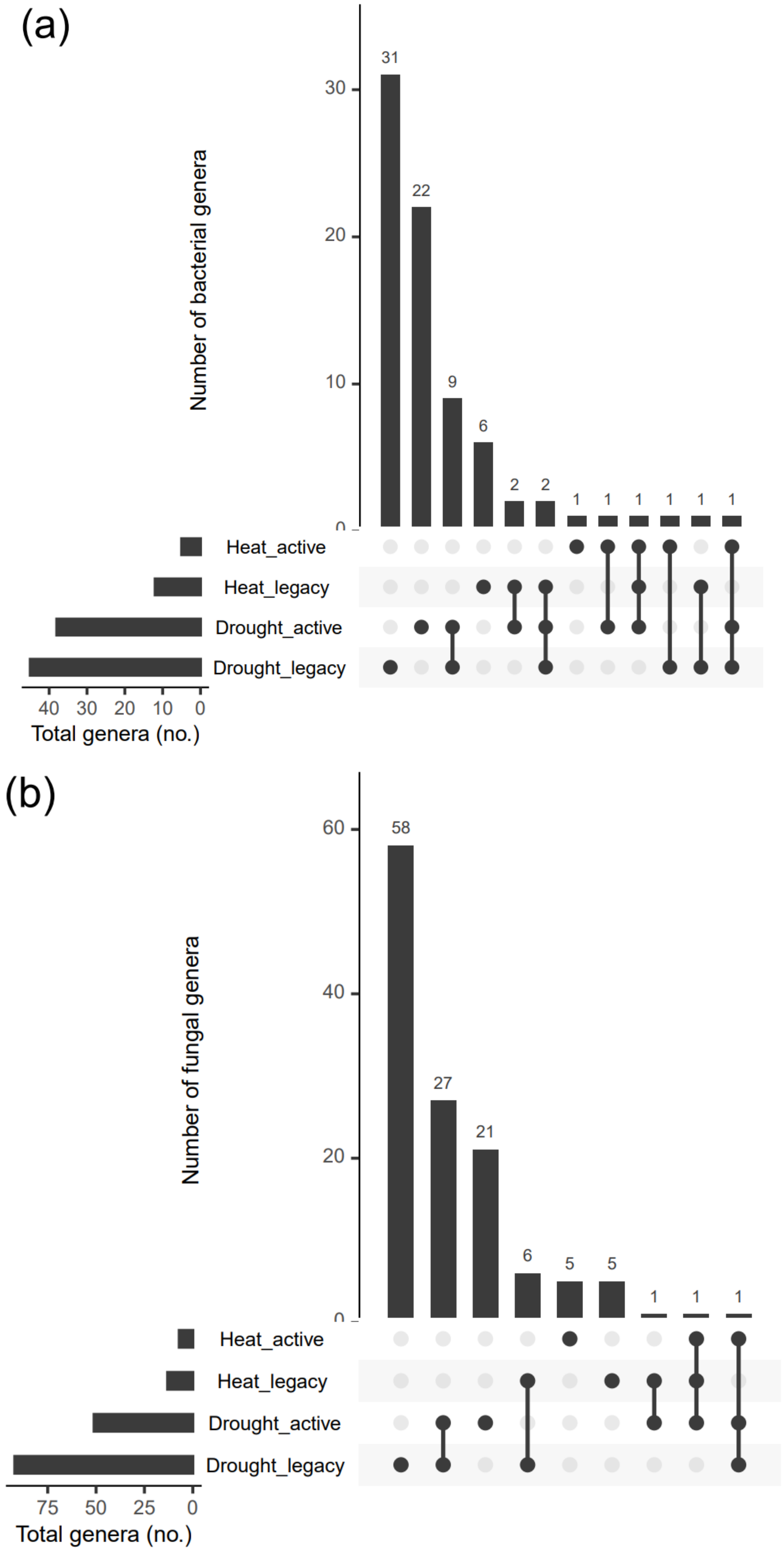


Figure S4. Overlap in bacterial (a) and fungal (b) genera significantly differentially abundant due to heat and drought in active treatment and recovery periods. Log fold change and significance was calculated using ANCOMBC.

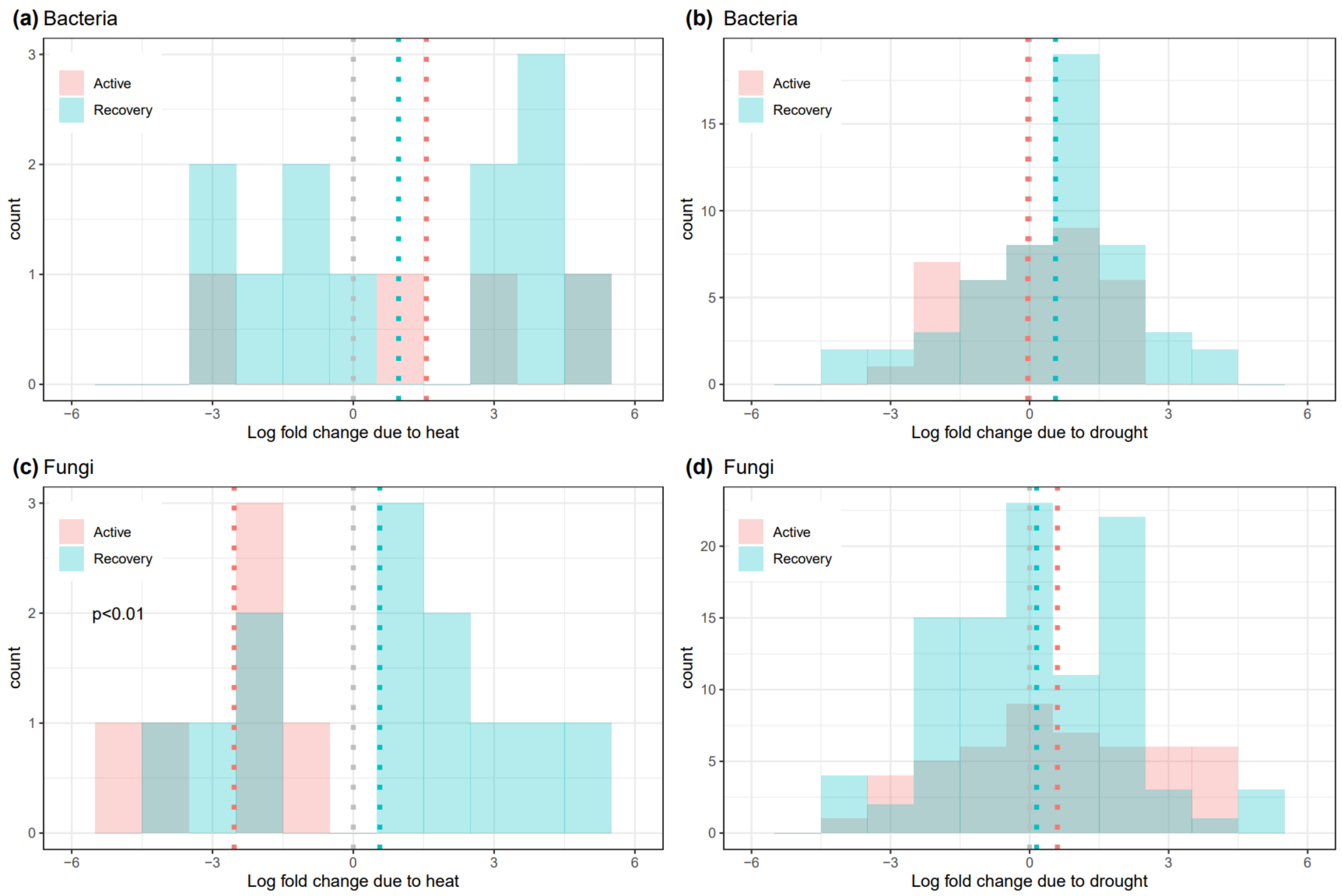


Figure S5. Histograms of significant log fold changes in recovery periods and active treatment. Log fold change and significance was calculated using ANCOMBC. Histograms are grouped by whether genera are bacteria or fungi, and climate factor. Colour indicates whether taxa were differentially abundant in active treatment (red) or recovery periods (blue). Grey dotted lines indicate 0, while red and blue dotted lines indicate the mean of the log fold change in active or recovery periods respectively. When significant, the *p*-value from student's t-tests comparing active and recovery response distributions is displayed on the plot.

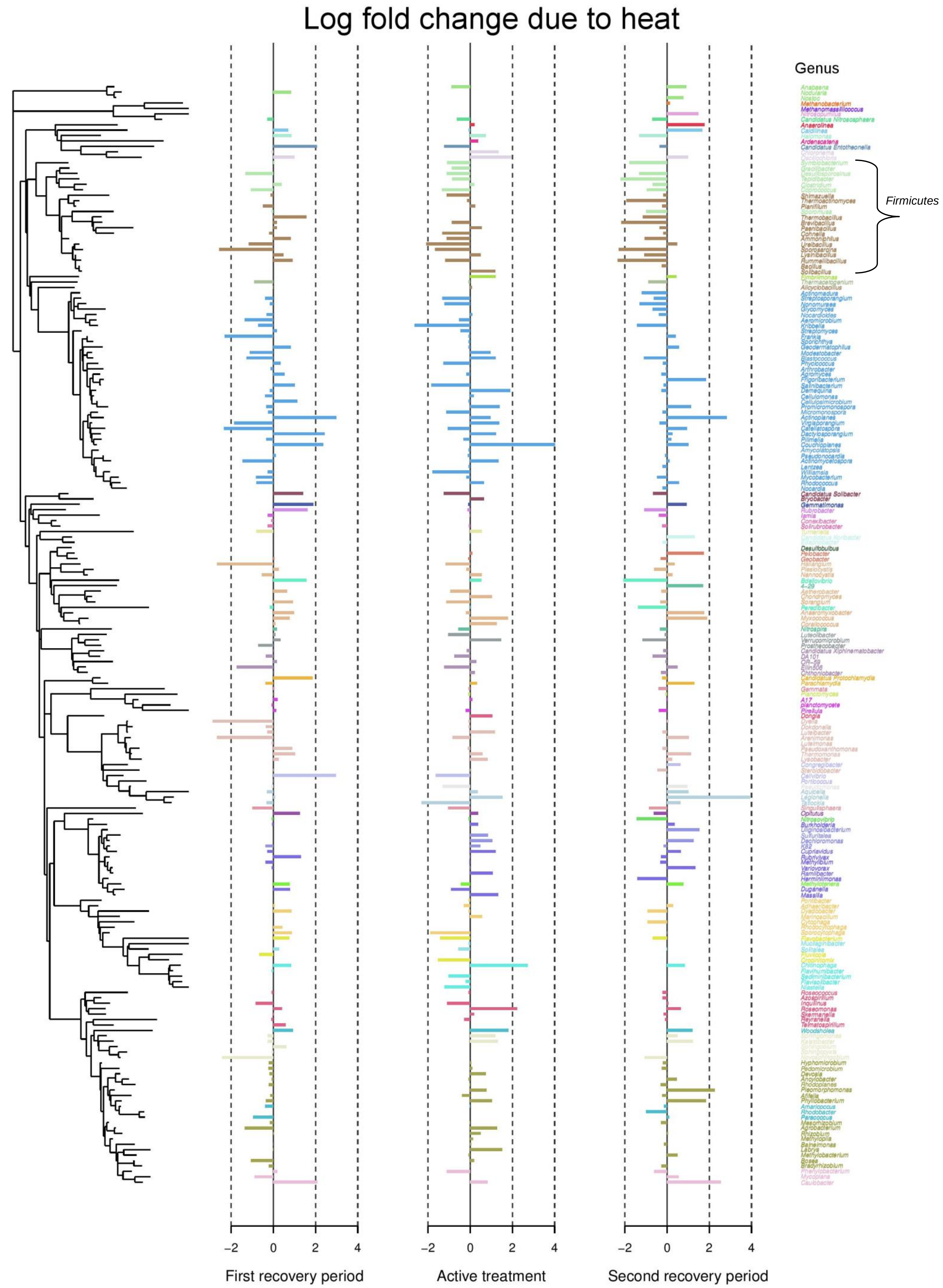


Figure S6. Differential abundance due to heat in bacteria, organized by phylogenetic relationship. Significant and non-significant differential abundances are included to detect signal. Log fold change was calculated using ANCOMBC. An example of a phylum with consistent responses to heat, *Firmicutes*, is indicated on the right.

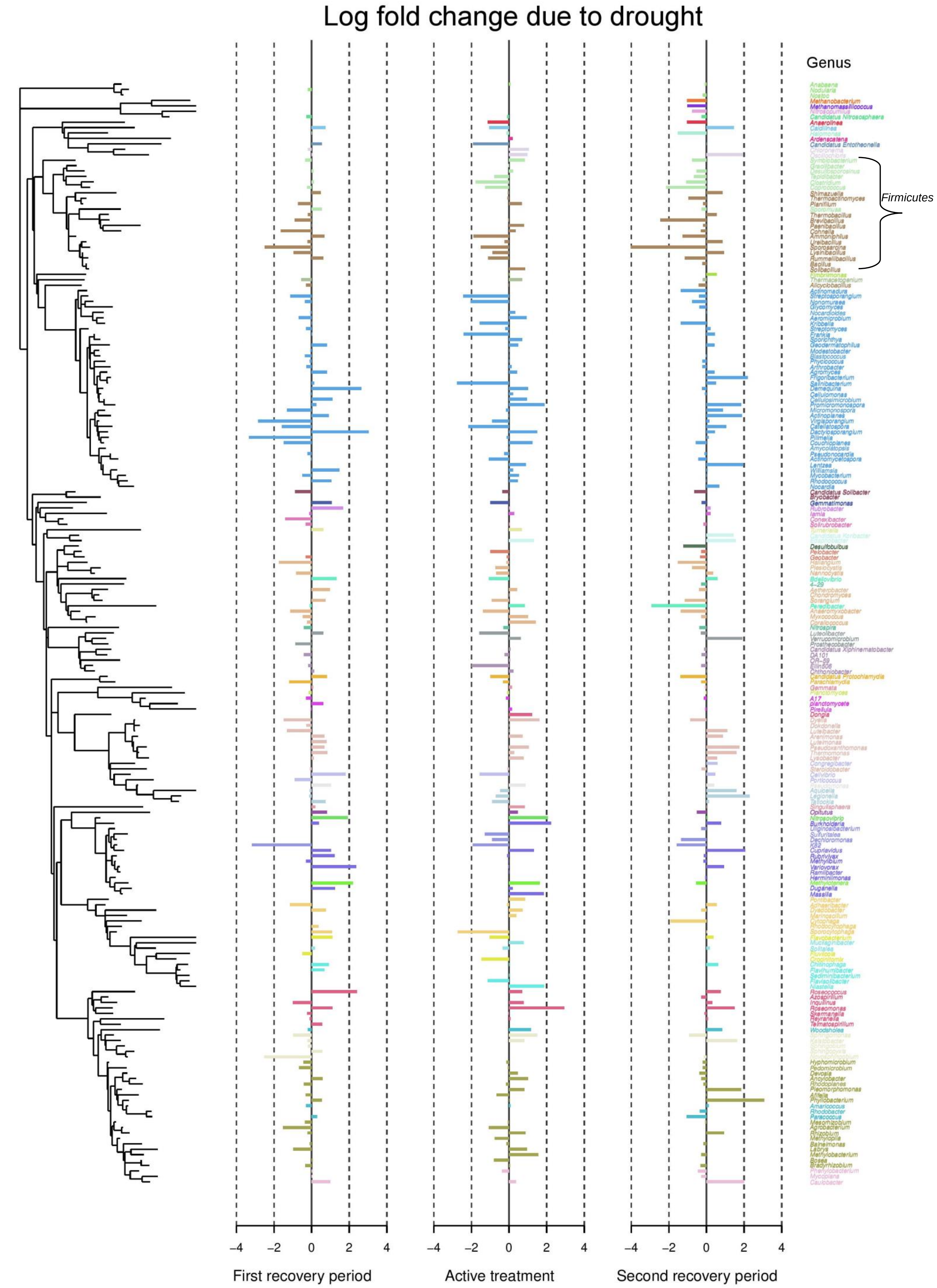


Figure S7. Differential abundance due to drought in bacteria, organized by phylogenetic relationship. Significant and non-significant differential abundances are included to detect signal. Log fold change was calculated using ANCOMBC. An example of a phylum with consistent responses to drought, *Firmicutes*, is indicated on the right.

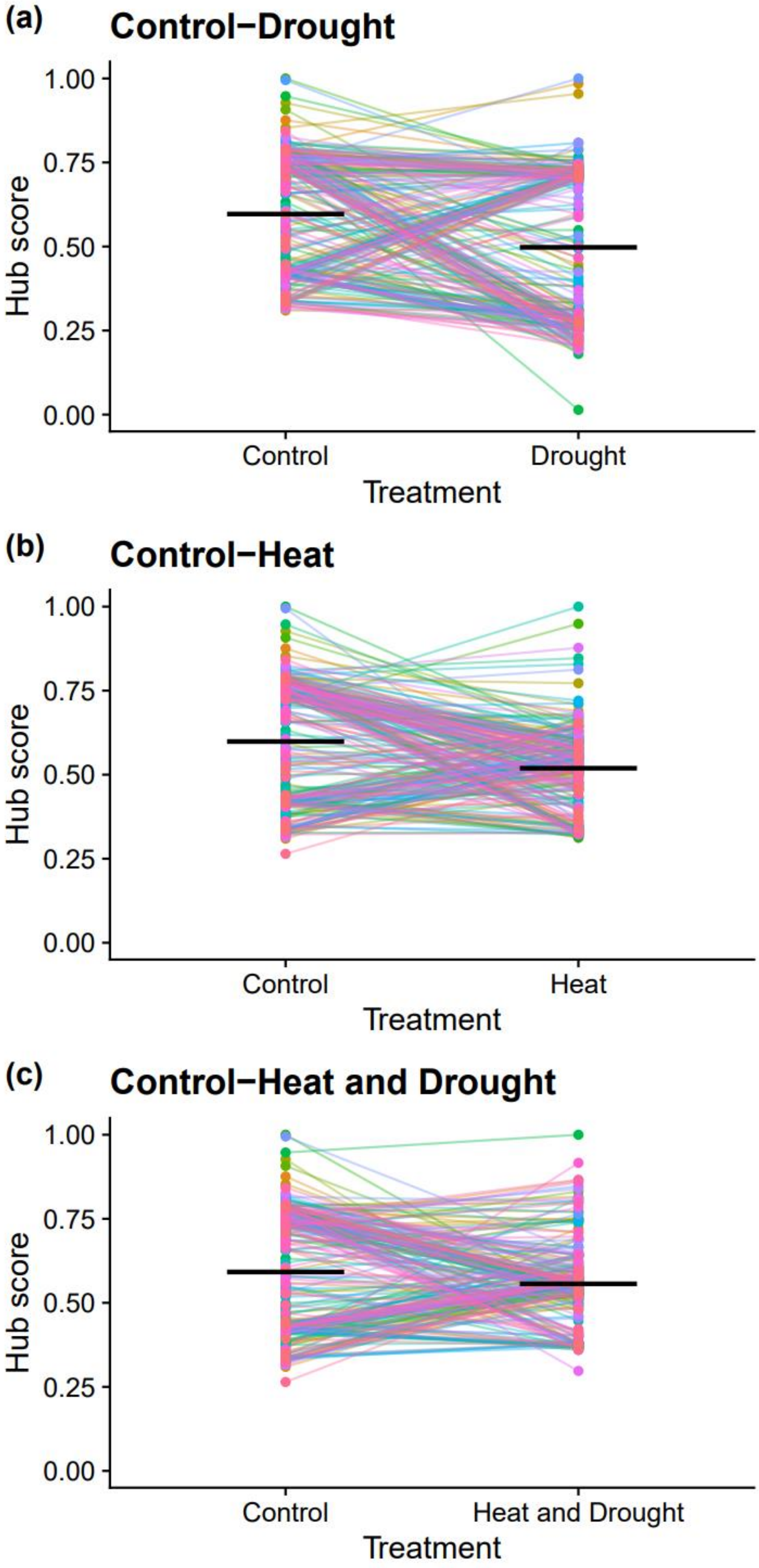


Figure S8. Hub scores of genera detected in both control plots and climate treated plots. Each point is a genus, and points belonging to the same genus are connected by a line across different treatments. Black bars indicate the mean hub score of the genera in that treatment. Scores tend to decrease overall in heat and drought, however there is considerable variation in how hub score is affected. Some genera remain the same, while other genera swap positions from high hub score to low hub score and vice versa.
